# Myosin 10 supports mitotic spindle bipolarity by promoting PCM integrity and supernumerary centrosome clustering

**DOI:** 10.1101/2022.02.08.479580

**Authors:** Yang-In Yim, Antonio Pedrosa, Xufeng Wu, Richard E. Cheney, John A. Hammer

## Abstract

Myosin 10 (Myo10) is a member of the MyTH4/FERM domain family of unconventional, actin-based motor proteins. Studies have implicated Myo10 in supporting cell adhesion via its integrin-binding FERM domain, and spindle positioning and spindle pole integrity via its microtubule-binding MyTH4 domain. Here we characterized Myo10’s contribution to mitosis using Myo10 knockout HeLa cells and MEFs isolated from a Myo10 knockout mouse. Most notably, both of these knockout cells exhibit a pronounced increase in the frequency of multipolar spindles. Staining of unsynchronized metaphase cells showed that the primary driver of spindle multipolarity in knockout MEFs and knockout HeLa cells lacking supernumerary centrosomes is PCM fragmentation, which creates y-tubulin-positive, centriole-negative microtubule asters that serve as additional spindle poles. For HeLa cells possessing supernumerary centrosomes, Myo10 depletion further accentuates spindle multipolarity by impairing centrosome clustering. These results argue, therefore, that Myo10 supports spindle bipolarity by maintaining PCM integrity in both normal and cancer cells, and by promoting supernumerary centrosome clustering in cancer cells. Finally, we present evidence that the defect in spindle pole integrity in Myo10 knockout cells is likely due to a defect in pole stability rather than pole maturation, and that Myo10 promotes supernumerary centrosome clustering at least in part by promoting cell adhesion during mitosis.

## Introduction

Myosin 10 (Myo10) is a member of the MyTH4/FERM domain family of unconventional myosins (reviewed in [1-5]). These actin-based motors are unusual in that they interact with microtubules and integrins via their MyTH4 and FERM domains, respectively. Myo10 is known as the “filopodial myosin” because it accumulates dramatically at the tips of filopodia [6-8], and because cells that over-express or lack Myo10 exhibit increases and decreases in filopodia number, respectively [6, 9-18]. Consistent with these observations, Myo10 strongly prefers to walk on bundled actin filaments like those that comprise filopodia [19-21], and it can be seen to move out filopodia at ∼800 nm/s [6, 22]. Perhaps most importantly for this study, Myo10 promotes the formation of adhesions within filopodia by virtue of its FERM domain-dependent binding to β1-integrin [23-25]. Filopodial adhesions sense extracellular matrix (ECM) stiffness and topography [26-29], generate traction force [9, 26, 28, 30-33], and often mature into focal adhesions upon cell advance [27, 29, 34-38]. Consistently, Myo10-dependent filopodial adhesions have been implicated in cell migration and cancer cell metastasis (reviewed in [9, 39, 40]).

While the bulk of cell biological studies to date have focused on Myo10’s role in the structure and function of filopodia, which are generally devoid of microtubules, Myo10 may also play important roles in microtubule-dependent processes via its microtubule-binding MyTH4 domain. The best evidence for this to date has come from studies focusing on Myo10’s role in meiosis and mitosis. First, Bement and colleagues showed that Myo10 localizes to the portion of the frog egg meiotic spindle in contact with the actin cortex, and that Myo10 depletion disrupts nuclear anchoring, spindle assembly and spindle-cortex association in eggs [41]. Subsequently, Bement and colleagues showed that Myo10 localizes to spindles and spindle poles in embryonic frog epithelial cells undergoing mitosis, and that Myo10 knockdown cells exhibit defects in spindle anchoring, spindle length, and metaphase progression [42]. They also showed that Myo10-depleted epithelial cells enter mitosis as bipolar but become multipolar during anaphase due to spindle pole fragmentation, arguing that Myo10 is required for spindle pole integrity [42]. Recent work from Pellman and colleagues using HeLa cells has shown that Myo10 is also required for spindle positioning [43]. Finally, Myo10 has been implicated along with the pole-focusing, microtubule minus end-directed kinesin 14 family member HSET in the microtubule-dependent clustering of supernumerary centrosomes based on RNAi screens for genes whose expression promotes such clustering [44].

Regarding Myo10’s contribution to spindle pole integrity, as proposed by Bement and colleagues [42], centrosomes begin a process starting in G2 called centrosome maturation that transforms them into the poles of the mitotic spindle (reviewed in [45, 46]). Maturation involves the recruitment of additional PCM components and microtubule-nucleating proteins like ⍰-TuRC, and results in a structure that differs from interphase centrosomes it being larger and more potent at nucleating microtubule assembly. These changes allow the spindle pole to support the assembly of the mitotic spindle and to resist the chromosomal and mitotic forces that are placed on it during mitosis. Consistently, defects in pole maturation lead to pole fragmentation during mitosis, and pole/PCM fragmentation is one of two defects that cause spindle multipolarity in cells lacking supernumerary centrosomes (the other being centriole disengagement; reviewed in [47]).

Four observations made by Bement and colleagues served to link the spindle pole fragmentation phenotype they saw in Myo10 morphants to a role for Myo10’s MyTH4 domain in promoting spindle pole integrity [42]. First, endogenous Myo10 was found concentrated at spindle poles. Second, Myo10’s localization overlapped significantly with that of TPX2, a microtubule binding protein and spindle assembly factor that is enriched at and near spindle poles, where it promotes branching microtubule nucleation [48-50]. Third, Myo10’s MyTH4/FERM domain was found to interact with TPX2 *in vitro*. Finally, Myo10 knockdown cells exhibited a significant reduction in the enrichment of TPX2 at and near poles. Together, these observations argued that Myo10 is required to maintain spindle pole integrity by recruiting TPX2, and that, without the stabilizing effect of TPX2, spindle poles in Myo10 morphants fragmented when subjected to the strong chromosomal and spindle forces that ramp up during metaphase.

With regard to Myo10’s role in spindle positioning in HeLa cells [43], Pellman and colleagues presented evidence that Myo10 localizes to subcortical actin clouds at the equator of dividing cells where it cooperates in a non-redundant and MyTH4 domain-dependent fashion with cortical dynein to position the mitotic spindle by engaging the tips of astral microtubules. That said, Toyoshima and colleagues [51] showed in an earlier study that the ability of integrin-based adhesions to orient the mitotic spindle parallel to the substratum requires Myo10. This result argued that Myo10 promotes spindle positioning at least in part by promoting adhesion to the ECM, presumably via its FERM domain-dependent interaction with integrins. Finally, whether Myo10 requires its microtubule-binding MyTH4 domain to promote supernumerary centrosome clustering is not known. In one reasonable scenario, Myo10 present in subcortical actin clouds at the cell equator could cooperate in a MyTH4 domain-dependent manner not only with cortical dynein to position the spindle, but also with HSET to promote supernumerary centrosome clustering [44].

Here we show that HeLa cells and MEFs lacking Myo10 exhibit a three-to five-fold increase in the frequency of multipolar spindles. By staining unsynchronized, metaphase cells for DNA, centrioles and y-tubulin, we show that the primary cause of spindle multipolarity in Myo10 knockout (KO) HeLa cells lacking supernumerary centrosomes and in Myo10 KO MEFs is PCM fragmentation, which creates y-tubulin-positive, centriole-negative microtubule asters that serve as additional spindle poles. While PCM fragmentation also contributes significantly to the formation of multipolar spindles in Myo10 KO HeLa cells that possess supernumerary centrosomes, the primary cause of spindle multipolarity in these cells is an inability to cluster their extra centrosomes. Together, these data show that Myo10 supports spindle bipolarity by maintaining PCM/spindle pole integrity in both normal and cancer cells, and by promoting supernumerary centrosome clustering in cancer cells. Finally, we present evidence that Myo10’s contribution to spindle pole integrity probably involves a role in maintaining pole stability rather than promoting pole maturation, and that its contribution to supernumerary centrosome clustering likely involves a role in promoting cell adhesion during mitosis.

## Results

### Myo10 KO HeLa cells exhibit a pronounced increase in the frequency of multipolar spindles

We used CRISPR-Cas9-based genome editing in an effort to abrogate Myo10 expression in HeLa cells, as these cells have been used extensively to explore Myo10 functions, including those related to mitosis [42, 43]. Of note, HeLa cells are polyploid (they contain 70-90 chromosomes instead of the 46 in normal human cells), and they appear to contain at least three Myo10 alleles based on analyses of chromosomal duplications [52]. We inserted guide sequences targeting Exon 3 of the Myo10 gene into plasmid pSpCas9(BB)-2A-GFP to create the construct pSpCas9(BB)-2A-GFP-ghM10-Exon3, which we then introduced into HeLa cells by electroporation. Two days later, GFP-positive cells were subjected to single-cell sorting into 96-well plates. This effort yielded only two putative Myo10 KO clones (KO-1 and KO-2) from 384 wells, or ∼0.5% of sorted cells. This low number likely reflects several factors, including the stresses incurred by single cell sorting (only ∼30% of control cells sorted to single cells grew up), the defects in mitosis that occur when Myo10 is depleted (see below), and the fact that these defects are more pronounced at lower cell densities (see below). Western blots of whole cell extracts prepared from WT HeLa, KO-1 and KO-2 and probed with an antibody to Myo10 showed a large reduction in Myo10 levels in KO-1 and KO-2 (Fig. 1A), with scans of replicate blots showing that these cells contain 7.1% and 13.8% of normal Myo10 levels, respectively (Fig. S1A). Sequencing of PCR products that span the portion of Exon 3 targeted by the guide sequences revealed two frameshift alleles and one missense allele in both KO-1 and KO-2 (DNS). We conclude, therefore, that both KO lines likely contain two nonsense alleles and one missense allele, making them hypomorphs for Myo10. While not complete KOs, we refer to these two lines as KO lines for the sake of simplicity.

**Figure 1.**
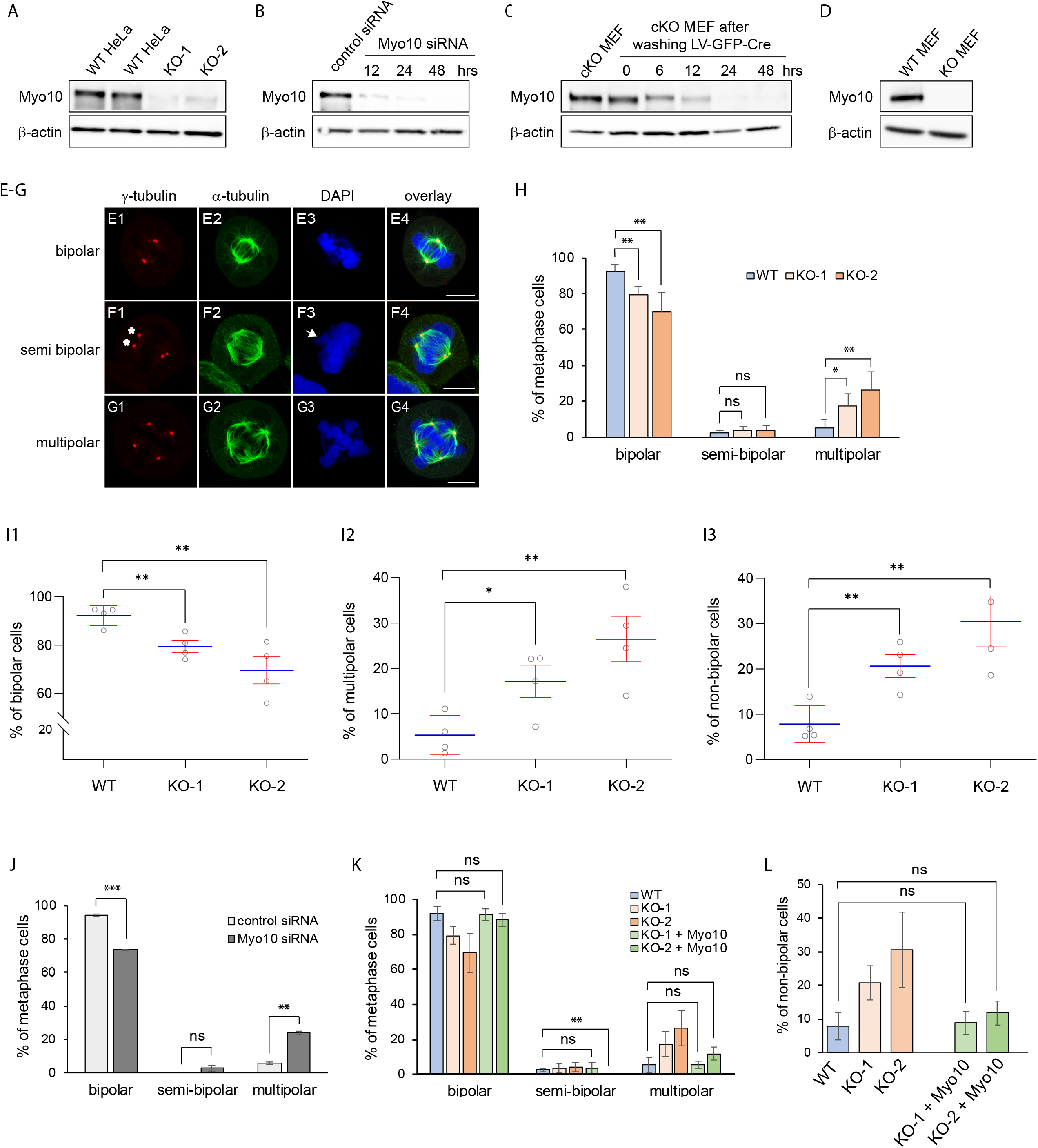
Myo10 depleted cells exhibit a significant increase in the frequency of multipolar spindles at metaphase. (A-D) Westerns blots of whole cell extracts prepared from WT and Myo10 KO HeLa cells (A), HeLa cells treated twice for 48 hrs each with a control, nontargeting siRNA or twice with a Myo10 Smartpool siRNA and collected 12, 24 and 48 hrs after starting the second treatment (B), non-transduced cKO MEFs and cKO MEFs 0, 6, 12, 24 and 48 hrs after transduction with lentiviral cre (C), and MEFs from a WT B6 mouse and from the tm1d Myo19 KO mouse (D) probed for Myo10 and β-actin as a loading control. (E1-E4) A representative example of a metaphase HeLa cell harboring a bipolar spindle stained for γ-tubulin, α-tubulin, and DNA (DAPI), along with the overlay (see also Z-Stack Movie 1). (F1-F4) Same as E1-E4 except that this HeLa cell harbored a semi-bipolar spindle (see also Z-Stack Movie 2). Two γ- tubulin-positive poles with asterisk indicate they were in different z planes, resulting in chromosome protrusion (arrow in F3). (G1-G4) Same as E1-E4 except that this HeLa cell harbored a multipolar spindle (see also Z-Stack Movie 3). (H) Quantitation at metaphase of the percent of bipolar, semi-bipolar, and multipolar spindles in WT HeLa and HeLa Myo10 KO lines KO-1 and KO-2 (281 WT cells, 289 KO-1 cells and 307 KO-2 cells from 4 experiments). (I1-I3) Comparison of WT Hela with KO-1 and KO-2 with regard to percent bipolar cells (I1), percent multipolar cells (I2) and percent non-bipolar cells (semi-bipolar plus multipolar), all at metaphase (calculated from the raw data in (H); shown as SEMs). (J) Quantitation at metaphase of the percent of bipolar, semi-bipolar, and multipolar spindles in HeLa cells treated twice (each for 48 hrs) with a control, nontargeting siRNA or with Myo10 Smartpool siRNA (173 control cells and 234 KD cells from 2 experiments). (K) Quantitation at metaphase of the percent of bipolar, semi-bipolar, and multipolar spindles in WT HeLa, KO-1, and KO-2 cells, and in KO-1 and KO-2 cells 24 hrs post-transfection with mScarlet-Myo10 (103 KO-1 cells and 120 KO-2 cells from 4 experiments; only cells expressing mScarlet-Myo10 were scored; only statistical values for WT versus rescued KO-1 and KO-2 cells are shown; the values for WT HeLa, KO-1 and KO-2 are from (H)). (L) Quantitation at metaphase of the percent of non-bipolar spindles (semi-bipolar plus multipolar) in WT HeLa, KO-1, and KO-2 cells, and in KO-1 and KO-2 cells 24 hrs post-transfection with mScarlet-Myo10 (only statistical values for WT versus rescued KO-1 and KO-2 cells are shown; the values for WT HeLa, KO-1 and KO-2 are from (H)). All the mag bars are 10 μm.

To identify possible defects in mitotic spindle organization upon Myo10 depletion, we fixed and stained unsynchronized WT HeLa, KO-1 and KO-2 for γ-tubulin to label spindle poles, α-tubulin to label spindle microtubules, and DAPI to label chromosomes. Cells identified as being at metaphase were optically sectioned in 0.25 μm intervals, and the sections used to count spindle pole numbers and to determine spindle and chromosome organization in 3D. Using these images, cells were scored as being either bipolar (this category included not only cells with just two spindle poles, one at each end of a normal-looking bipolar spindle, but also pseudo-bipolar cells where one or both poles at each end of a normal looking bipolar spindle were created by the clustering of supernumerary centrosomes; Fig. 1, E1-E4; Z-Stack Movie 1), semi-polar (this category contained cells possessing two major poles and a normal looking spindle, but also a minor pole in a different focal plane that contributed some microtubules to the spindle; Fig. 1, F1-F4; Z-Stack Movie 2), or multipolar (this category contained cells with more than two major spindle poles; Fig. 1, G1-G4; Z-Stack Movie 3). Scoring of >300 cells per line over four independent experiments showed that KO-1 and KO-2 exhibit 11.8% and 22.7% decreases in the frequency of bipolar spindles, respectively (Fig. IH and I1), 3.2-fold and 5.0-fold increases in the frequency of multipolar spindles (Fig. 1H and I2), and 2.6-fold and 3.9-fold increases in the frequency of non-bipolar spindles (semi-polar plus multipolar) (Fig. 1H and I3). We conclude, therefore, that HeLa cells with reduced levels of Myo10 exhibit a pronounced increase in the frequency of multipolar spindles.

### Myo10 KO HeLa cells exhibit an increase in the frequency of spindles that are not parallel to the substratum

For non-polarized cells dividing in 2D, the balance of forces driving spindle positioning are such that the spindle’s long axis is usually parallel to the substratum, resulting in the two poles being about equidistant from the substratum i.e. in the same Z-plane. Optical sectioning of ∼200 metaphase cells exhibiting just two poles showed that the average distance in Z separating the two spindle poles was significantly greater in both KO lines than in WT HeLa (Fig. 2A; 2D shows the distributions of these distances in 1 μm intervals; see also Z-Stack Movie 4). This phenotype might reflect a defect in the adhesion of Myo10 KO cells, resulting in an imbalance in the pulling forces driving spindle positioning ([51]; see Discussion). While the mean values for the separation of the two poles in Z was also elevated in both KO lines at anaphase and telophase (Fig. 2B, 2C; Z-Stack Movies 5 and 6), the differences were not quite statistically significant at anaphase and far from significant at telophase. This result indicates that the defect in positioning the metaphase spindle relative to the substratum seen when Myo10 is depleted is largely corrected as mitosis progresses to telophase.

**Figure 2.**
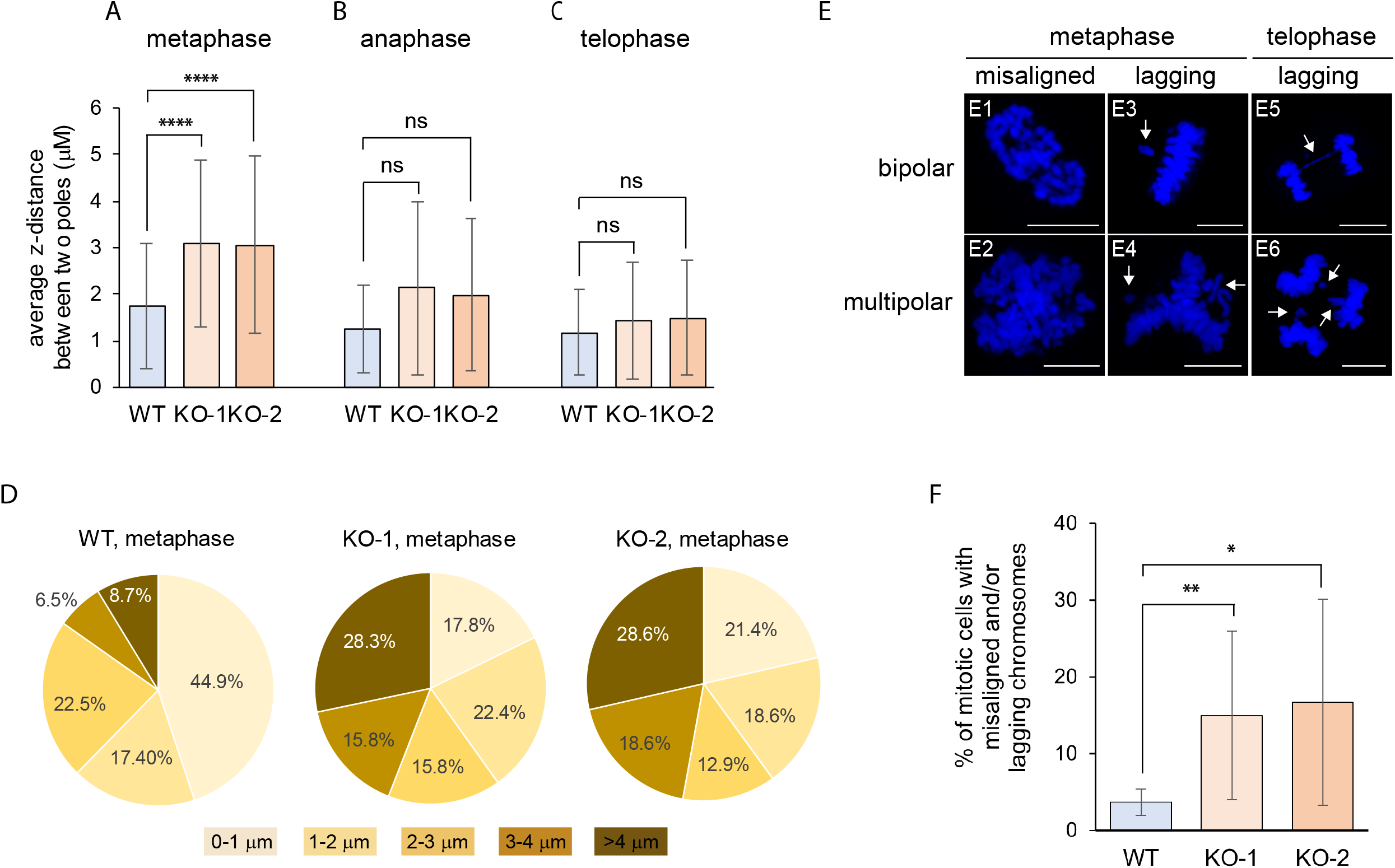
Myo10 KO cells at metaphase exhibit increased frequencies of spindles that are not parallel to the substratum and increased frequencies of misaligned and lagging chromosomes. (A-C) Average distance in Z between the two poles of bipolar WT HeLa, KO-1 and KO-2 at metaphase (A), anaphase (B) and telophase (C) (180 WT cells, 234 KO-1 cells and 228 KO-2 cells from 4 experiments; see also Z-Stack Movies 4-6). (D) The distributions of the values in (A) in 1 um intervals. (E1-E6) Shown are representative examples of misaligned chromosomes at metaphase in KO-1 cells undergoing bipolar mitosis (E1) or multipolar mitosis (E2), and lagging chromosomes at metaphase and telophase in KO-1 cells undergoing bipolar mitosis (E3 and E5, respectively) or multipolar mitosis (E4 and E6, respectively) (arrows point to lagging chromosomes). (F) Percent of mitotic WT HeLa, KO-1 and KO-2 that possess misaligned and/or lagging chromosomes (258 WT cells, 308 KO-1 cells and 330 KO-2 cells from 4 experiments). All the mag bars are 10 μm.

### Myo10 KO HeLa cells exhibit increased frequencies of chromosome misalignment and lagging chromosomes

Cells with multipolar or improperly positioned spindles often exhibit chromosome misalignment at metaphase and lagging chromosomes later in mitosis due to merotelic or missing kinetochore attachments. Imaging of mitotic cells following fixation and staining for DNA, microtubules and spindle poles revealed evidence between metaphase and telophase of these chromosome distribution defects in Myo10 KO cells undergoing both bipolar and multipolar divisions (Fig. 2, E1-E6; the arrows in E3-E6 indicate lagging chromosomes). Indeed, scoring showed that the percentage of mitotic cells exhibiting chromosome misalignment and/or lagging chromosomes was 4.0-and 4.5-fold higher in KO-1 and KO-2, respectively, than in WT HeLa cells (Fig. 2F). This result is consistent with the defects in spindle bipolarity and spindle positioning exhibited by these KO lines.

### HeLa cells in which Myo10 levels were reduced using siRNA-mediated knockdown also exhibit an increase in the frequency of multipolar spindles

KO-1 and KO-2 initially grew very slowly, only approaching an approximately normal growth rate about two months in culture (at which point we started collecting data). This behavior suggests that these clonal isolates underwent some kind of genetic or epigenetic change after CRISPR-based genome editing that ameliorated to some extent the cellular defects responsible for their slow growth rate. Given this, and given that both of our KO clones are hypomorphs, we decided to also examine HeLa cells in which Myo10 was transiently depleted using siRNA-mediated knockdown (KD). Relative to control cells that received a control, nontargeting siRNA, cells that received two rounds of treatment with Myo10 SmartPool siRNA exhibited near complete depletion of Myo10 48 hours after the second round of treatment (Fig. 1B and Fig. S1B). At 48 hours post-transfection, unsynchronized, metaphase control and KD cells were scored for spindle phenotype as described above for WT, KO-1 and KO-2 cells. Consistent with KO-1 and KO-2 cells, Myo10 KD cells exhibited a 20.7% reduction in the frequency of bipolar spindles and a 4.1-fold increase in the frequency of multipolar spindles (Fig. 1J). We conclude, therefore, that this core Myo10 KO phenotype is replicated in Hela cells where Myo10 is transiently depleted using RNAi.

### MEFs isolated from a Myo10 conditional KO mouse and transduced with lenti-Cre phenocopy Myo10 KO HeLa cells

We recently described the creation of a Myo10 conditional knockout (cKO) mouse (tm1c) and several phenotypes exhibited by this mouse following its cross with a global Cre-deleter strain [13]. To provide additional support for the results obtained above using Myo10 KO HeLa cells, we treated primary MEFs isolated from this Myo10 cKO mouse with lentivirus expressing GFP-tagged Cre recombinase. Western blots showed that Myo10 was essentially undetectable 24 to 48 hours after lenti-cre transduction (Fig. 1C and Fig. S1C). Given this, non-transduced cKO MEFs and cKO MEFs 48 hours after transduction with lenti-Cre were scored for their metaphase spindle phenotype as described above for WT, KO-1 and KO-2 cells. As with Myo10 KO HeLa cells, scoring of >300 cells per line over three independent experiments showed that the lenti-Cre-transduced cKO MEFs exhibit a 35.6% decrease in the frequency of bipolar spindles, a 4.7-fold increase in the frequency of semipolar spindles, a 2.9-fold increase in the frequency of multipolar spindles, and a 3.6-fold increase in the frequency of non-bipolar spindles (semi-polar plus multipolar) (Fig. 3A). Also like Myo10 KO HeLa cells, optical sectioning of ∼200 metaphase cells exhibiting just two poles showed that the average distance separating the two spindle poles in Z was significantly greater in the lenti-Cre-transduced cKO MEFs than in the non-transduced cKO MEFs (Fig. 3B; 3C shows the distributions of these distances in 1 μm intervals). Finally, like Myo10 KO HeLa cells, the percentage of lenti-Cre-transduced cKO MEFs exhibiting chromosome misalignment and/or lagging chromosomes was 6.9-fold higher than for non-transduced cKO MEFs (Fig. 3D). We conclude, therefore, that cKO MEFs subjected to the conditional and rapid depletion of Myo10 largely phenocopy Myo10 KO HeLa cells as regards increased spindle multipolarity, abnormal spindle orientation relative to the substratum, and increased frequencies of chromosome misalignment and lagging chromosomes between metaphase and telophase.

**Figure 3.**
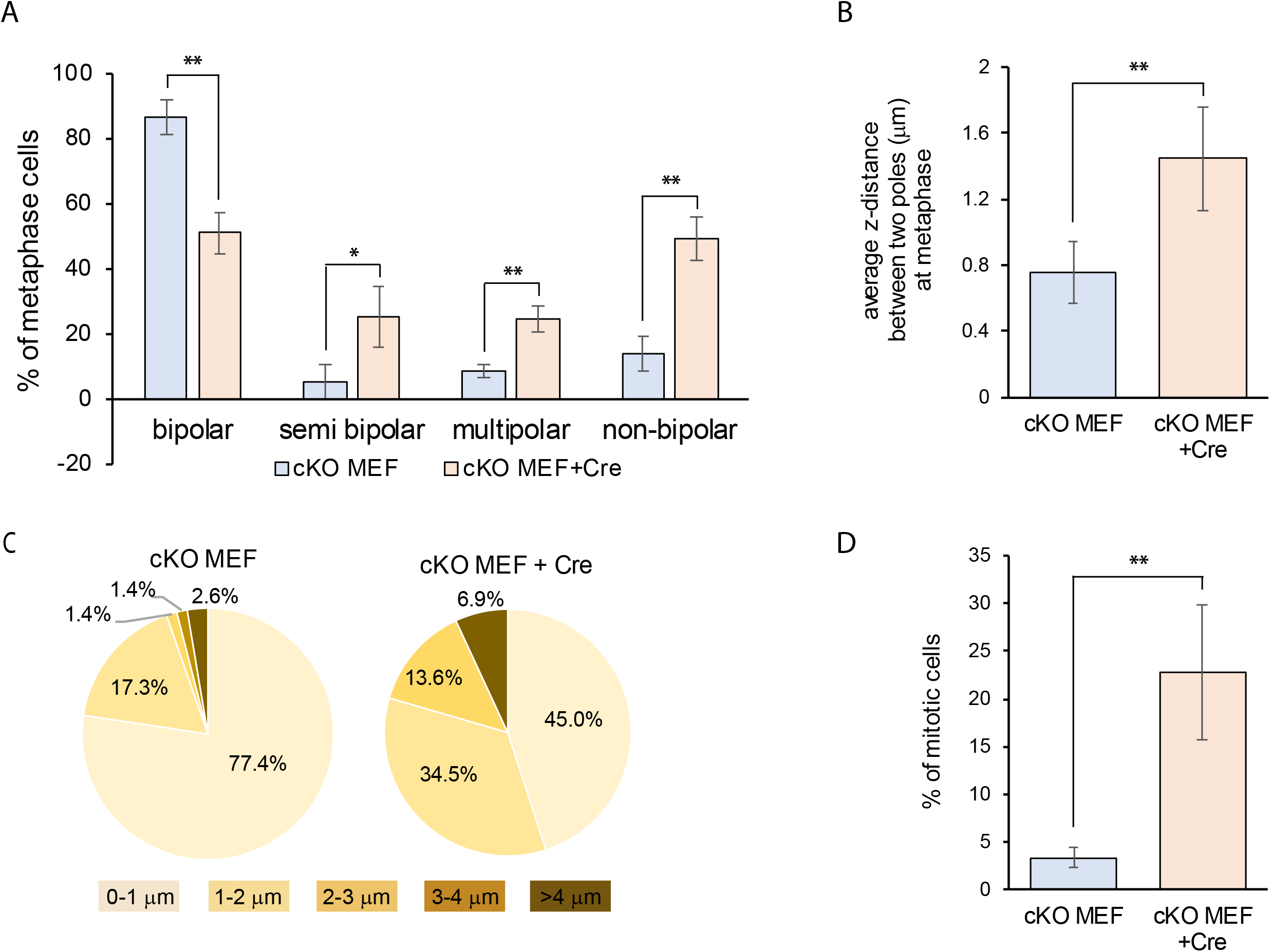
cKO MEFs transduced with lenti-Cre phenocopy Myo10 KO HeLa cells. (A) Quantitation at metaphase of the percent of bipolar, semi-bipolar, multipolar, and non-bipolar spindles in non-transduced cKO MEFs and cKO MEFs 48 hrs after lenti-cre transduction (159 cKO MEFS and 153 cKO MEFS plus Cre from 3 experiments). (B) Average distance in Z between the two poles of non-transduced cKO MEFs and cKO MEFs 48 hrs after lenti-cre transduction at metaphase (191 cKO MEFs and 208 cKO MEFs plus Cre over 3 experiments). (C) The distributions of the values in (B) in 1 um intervals. (D) Percent of mitotic, non-transduced cKO MEFs and cKO MEFs 48 hrs after lenti-cre transduction that possess misaligned and/or lagging chromosomes (169 cKO MEFs and 168 cKO MEFs plus Cre from 3 experiments).

### Myo10 KO MEFs from the straight Myo10 KO mouse exhibit an increase in the frequency of multipolar spindles whose magnitude depends on the phenotype of the embryo from which they are isolated and the density at which they are cultured

cKO MEFs following lenti-cre treatment exhibited several abnormalities in addition to those related to mitosis, including Golgi fragmentation, autophagy induction and p53 activation (data not shown). Given this, we decided to characterize the mitotic phenotype of MEFs isolated from the straight Myo10 KO mouse (tm1d; created by crossing the cKO Myo10 KO mouse (tm1c) with a global cre deleter strain; [13]). As described previously [13], the fate of embryos in single litters from the straight Myo10 KO mouse range from early embryonic lethality associated with exencephaly (a severe defect in neural tube closure) to viable mice exhibiting small body size, webbed digits, white belly spots, and microphthalmia. Given that the tm1d mouse is on a pure B6 background, this wide variation in embryo fate must be due to incomplete penetrance/variable expressivity rather than to variations between embryos in the inheritance of genetic modifiers [53] (although the non-genetic variations that dictate the fate of Myo10 KO embryos are unknown). We decided, therefore, to characterize MEFs isolated from both non-exencephalic and exencephalic tm1d embryos. We also decided to score the mitotic defects exhibited by these MEFs 24 hours after seeding them densely (5 × 10^4^ cells/cm^2^), at moderate density (2.5 × 10^4^ cells/cm^2^), and sparsely (0.5 × 10^4^ cells/cm^2^). This decision was based on measurements of growth rates for cells seeded at these three densities (referred to below and in the figures as “dense or D”, “moderate or M”, and “sparse or S”), where the growth of WT MEFs was unaffected by seeding density, while the growth of both non-exencephalic and exencephalic Myo10 KO MEFs was significantly slower at the sparse seeding density (Fig. 4A), as well as significantly slower than WT MEFs at the sparse seeding density (Fig. 4B). This result raised the possibility that Myo10 KO MEFs exhibit higher frequencies of mitotic defects and subsequent aneuploidy/cell senescence when provided with fewer spatial cues from neighboring cells.

**Figure 4.**
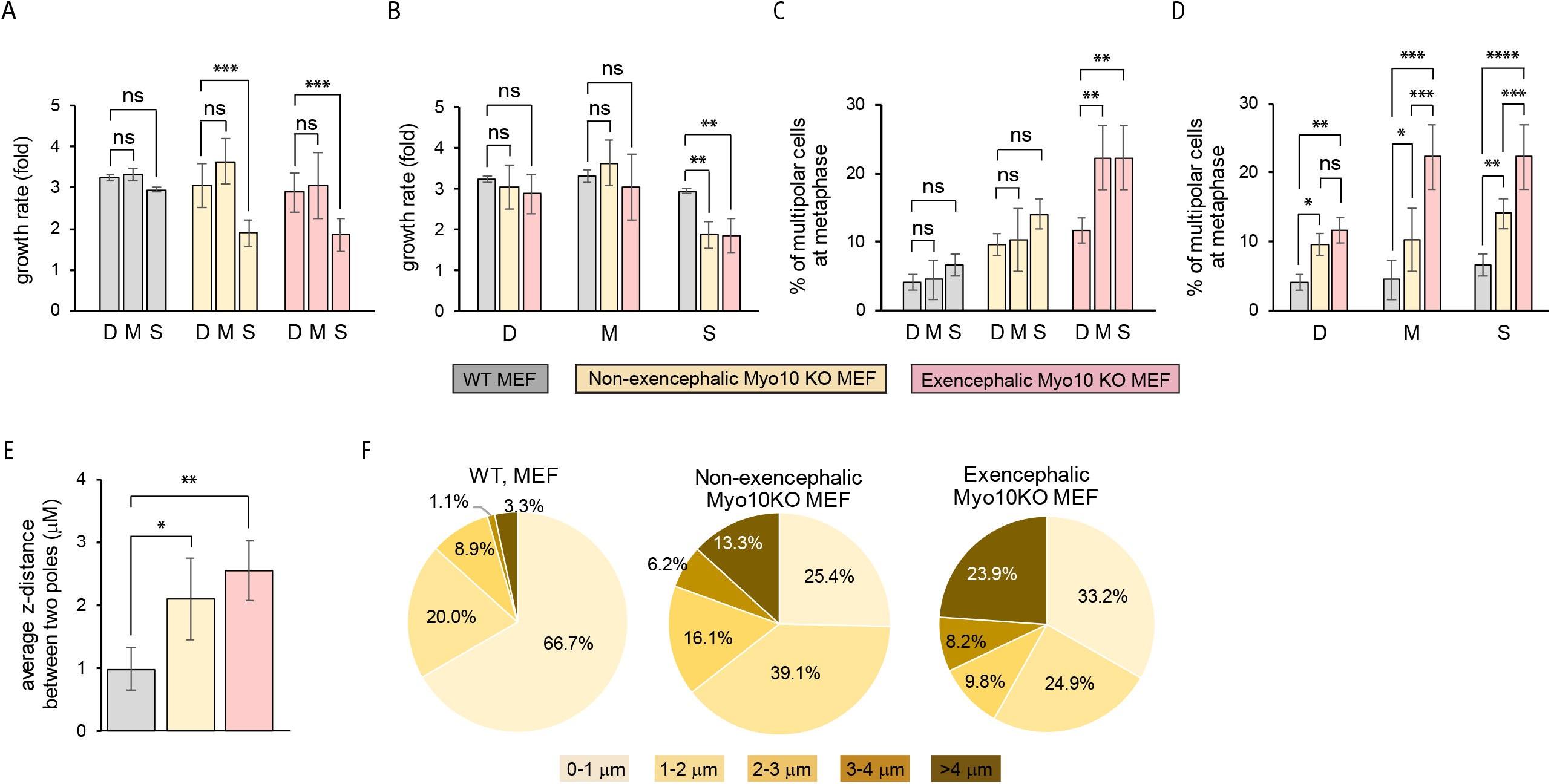
Myo10 KO MEFs exhibit spindle defects whose severity depends upon embryo source and culture density. (A) Growth rates of WT MEFs, MEFs isolated from non-exencephalic Myo10 KO embryos, and MEFs isolated from exencephalic Myo10 KO embryos and seeded at different cell densities (D, M and S stand for dense, moderate and sparse seeding, respectively; from 3 experiments, each done in duplicate). (B) The same data as in (A) but presented so as to allow comparison of the three types of MEF at each cell density. (C) Quantitation at metaphase of the percent of multipolar spindles in WT MEFs, non-exencephalic Myo10 KO MEFs, and exencephalic Myo10 KO MEFs grown at different cell densities (D: 147 WT MEFs, 145 non-exencephalic KO MEFs, and 186 exencephalic KO MEFs; M: 142 WT MEFs, 177 non-exencephalic KO MEFs, and 241 exencephalic KO MEFs; S: 161 WT MEFs, 209 non-exencephalic KO MEFs, and 257 exencephalic KO MEFs; from 4 experiments). (D) The same data as in (C) but presented so as to allow comparison of the three types of MEF at each cell density. (E) Average distance in Z at metaphase between the two poles of WT MEFs, non-exencephalic Myo10 KO MEFs, and exencephalic Myo10 KO seeded at moderate density. (F) The distributions of the values in (E) in 1 μm intervals.

To look for evidence that growth at lower cell densities does in fact increase the frequency of mitotic errors in Myo10 KO cells, we scored the frequency of multipolar spindles at metaphase in WT MEFs, non-exencephalic Myo10 KO MEFs, and exencephalic Myo10 KO MEFs 24 hours after seeding them at the three cell densities. While WT MEFs and non-exencephalic Myo10 KO MEFs were unaffected by cell density, exencephalic Myo10 KO MEFs exhibited higher frequencies of multipolar spindles as the cell density was lowered (Fig. 4C). Moreover, the frequencies for both KO MEFs were significantly higher than the frequency for WT MEFs at both moderate and sparse seeding densities (Fig. 4D). Additionally, the frequency exhibited by exencephalic Myo10 KO MEFs was significantly higher than the frequency exhibited by non-exencephalic Myo10 KO MEFs at both moderate and sparse seeding densities (Fig. 4D). Together, these results show that MEFs isolated from the straight Myo10 KO mouse exhibit a significant increase in the frequency of multipolar spindles, and that the severity of this defect depends on the phenotype of the embryo from which the MEFs are isolated, and on the density at which the MEFs are cultured. Finally, both non-exencephalic and exencephalic Myo10 KO MEFs grown at moderate density exhibit an increase in the frequency of spindles that are not parallel to the substate (Fig. 4E and 4F). These results provide further support for a defect in cell adhesion during mitosis when Myo10 is missing.

### The multipolar phenotype exhibited by Myo10 KO HeLa cells is rescued by re-expression of Myo10

The fact that the multipolar spindle phenotype exhibited by Myo10 KO HeLa cells is also seen in Myo10 KD HeLa cells, Myo10 cKO MEFs after cre expression, and straight Myo10 KO MEFs argues that it is indeed a consequence of Myo10 depletion/loss. That said, we thought it was important to show that the re-introduction of Myo10 into Myo10 KO HeLa cells rescues the multipolar spindle phenotype before proceeding with efforts to define the mechanism(s) underlying this phenotype. Towards that end, we transfected KO-1 and KO-2 with mScarlet-Myo10 and scored mScarlet-Myo10-expressing cells at metaphase for spindle phenotype 24 hours post-transfection. Statistical analyses showed that Myo10 re-expression restored the values for the percentage of metaphase cells with bipolar and multipolar spindles in both KO lines to WT HeLa levels (Fig. 1K). Moreover, combining the data for semi-bipolar and multipolar spindles into non-polar spindles showed that both KO lines were completely rescued by M10 re-expression (Fig. 1L). Together, these results paved the way for subsequent efforts described below to define the mechanism(s) underlying the multipolar phenotype.

### mScarlet-Myo10 localizes in mitotic HeLa cells to sites of cell adhesion and, to a much lesser extent, at/near spindle poles

The rescue experiments described above also provided information on Myo10’s localization in both interphase and mitotic cells. Consistent with images presented in numerous publications supporting Myo10’s designation as the “filopodial myosin” (reviewed in [1-5, 39, 40]), mScarlet-Myo10 localized dramatically to the tips of filopodia in rescued KO-1 cells during interphase (Fig. 5A1-A6). Also consistent with previously published images [43], mScarlet-Myo10 localized dramatically to the tips of retraction fibers in rescued KO-1 cells at metaphase (Fig. 5B1-B4). The localization of Myo10 as these actin-rich mitotic structures, which have been implicated for decades in the adhesion of mitotic cells [54-58], together with Myo10’s ability to link integrins to actin via its FERM domain and motor domain, respectively, suggests that Myo10 may play a significant role in driving adhesion during mitosis. Through focusing of metaphase cells showed that Myo10 is also present at the tips of dorsal filopodia, as observed previously [10] (Fig. 5C1-C4; Z-Stack Movie 7). Finally, in partial agreement with the results of Woolner et al [42] and Sandquist et al [59, 60], mScarlet-Myo10 appeared enriched to some degree at/near spindle poles and in the spindle in a subset of metaphase cells stained with Phalloidin and DAPI (Fig. 5D1 and D2; Z-Stack Movie 8). To further clarify the localization of mScarlet-Myo10 at/near spindle poles, we imaged metaphase cells stained for γ-tubulin. In a very small subset of cells (5 out of 171 metaphase cells imaged), we observed clear accumulation of mScarlet-Myo10 at or very near γ-tubulin-positive spindle poles (Fig. 5E1-E3; Z-Stack Movie 9), and in a very few cases in the spindle as well (Fig. 5F1-F3; Z-Stack Movie 10). In summary, then, the location of mScarlet-Myo10 in rescued KO-1 cells matched in large part the localization data reported previously for Myo10 in both interphase and mitotic cells (although clear localization at spindles and spindle poles was rare). Moreover, Myo10’s dramatic localization in retraction fibers suggests that it might promote supernumerary centrosome clustering at least in part by promoting cell adhesion during mitosis (see Discussion).

**Figure 5.**
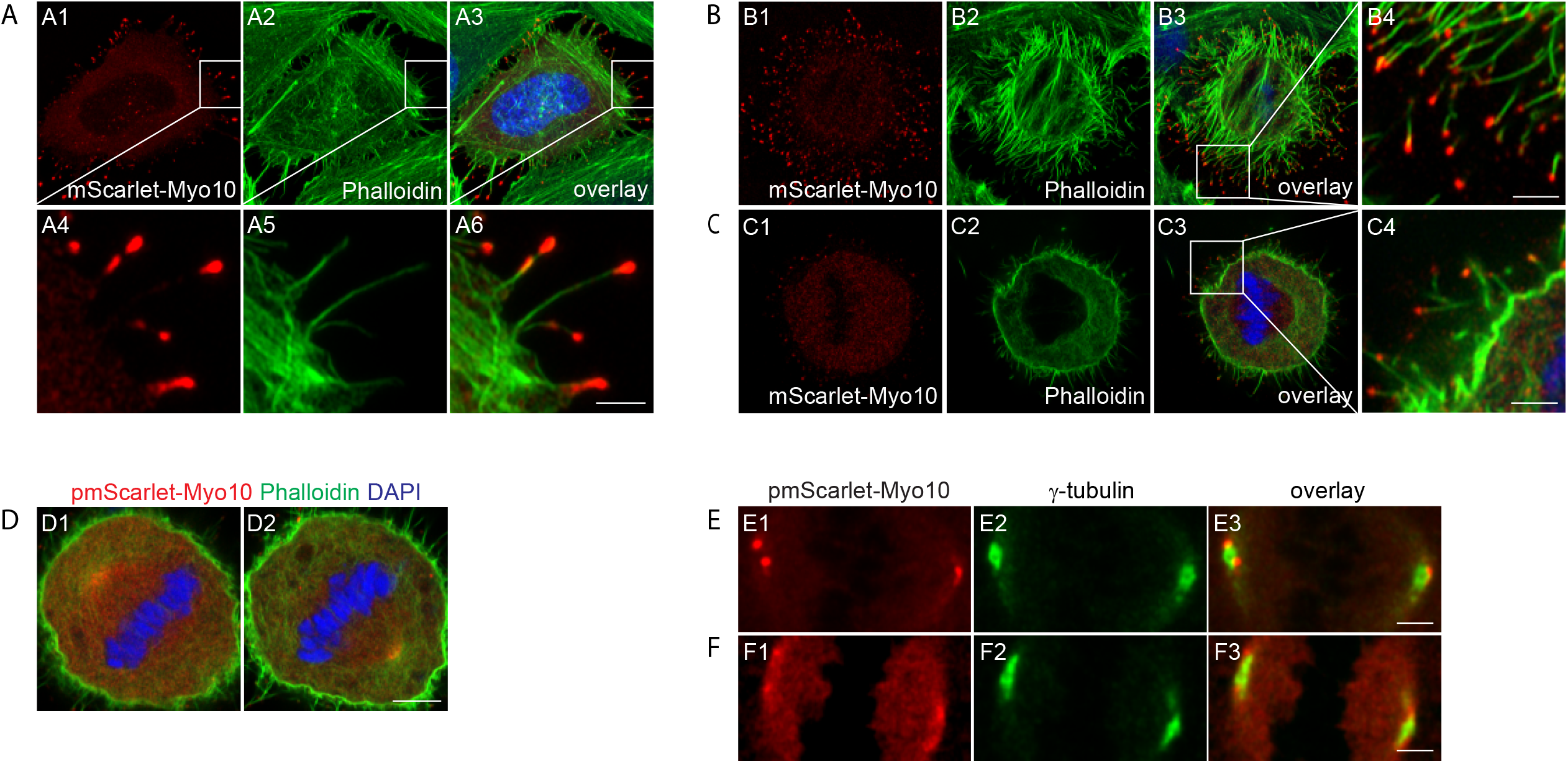
mScarlet-Myo10 localizes in mitotic HeLa cells to sites of cell adhesion and, to a much lesser extent, at/near spindle poles. (A1-A6) Representative example of an interphase Myo10 KO-1 cell expressing mScarlet-Myo10 that was fixed and stained with phalloidin, and imaged to show its ventral, Myo10-positive filopodia. (B1-B4) Representative example of a metaphase Myo10 KO-1 cell expressing mScarlet-Myo10 that was fixed and stained with phalloidin, and imaged to show its ventral, Myo10-positive retraction fibers. (C1-C4) The same cell as in (B) but imaged to show its dorsal, Myo10-positive filopodia (see also Z-Stack Movie 7). (D1 and D2) Example of a metaphase Myo10 KO-1 cell expressing mScarlet-Myo10 that was fixed and stained with phalloidin and DAPI, and imaged in two focal planes near the cell equator to show the modest amount of Myo10 that was sometimes present at/near spindle poles and in the spindle (see also Z-Stack Movie 8). (E1-E3, F1-F3) Examples of metaphase Myo10 KO-1 cells expressing mScarlet-Myo10 that were fixed and stained with for γ-tubulin, and imaged to show the modest amounts of Myo10 that were sometimes present at/near spindle poles (E1-E3; see also Z-Stack Movie 9) and, in some cases, the spindle as well (F1-F3; see also Z-Stack Movie 10). The mag bars in A6, B4, C4, E3 and F3 are 2 μm, and the mag bar in D2 is 5 μm.

### Multipolar spindles in Myo10 KO HeLa cells arise from a combination of PCM fragmentation and an inability to cluster supernumerary centrosomes

We used our Myo10 KO HeLa cells to search for the cause(s) of spindle multipolarity when Myo10 is depleted. As mentioned in the Introduction, four distinct cellular defects can give rise to multipolar spindles (reviewed in [47]). Two of these defects, cytokinesis failure and centriole overduplication, can give rise to multipolar spindles because they generate cells with extra centrosomes. The third defect, centriole disengagement, can give rise to multipolar spindles because both separated centrioles can serve as MTOCs. Finally, PCM fragmentation can give rise to multipolar spindles because the acentriolar PCM fragments that are generated can serve as MTOCs. Consistent with the results of Kwon et al [44] using Myo10 KD HeLa cells, we did not see a significant difference between WT and Myo10 KO HeLa cells in the average number of nuclei per cell (Fig. S2A1, S2A2), arguing that cytokinesis failure does not contribute significantly to their multipolar spindle phenotype. Centriole overduplication also does not appear to be driving the multipolar spindle phenotype, as unsynchronized WT and Myo10 KO HeLa cells have the same number of centrioles (Fig. S2B). For normal cells, which possess one centrosome, that would leave centriole disengagement and PCM fragmentation as the remaining possible causes of spindle multipolarity. For cancer cells, which often possess supernumerary centrosomes, the inability to cluster these extra centrosomes represents a third possible cause of spindle multipolarity (reviewed in [61-63]). Given that a large fraction of HeLa cells possesses supernumerary centrosomes (see below), and that Myo10 has been implicated in promoting supernumerary centrosome clustering to support spindle bipolarity [44], subsequent experiments were designed to distinguish between centriole disengagement, PCM fragmentation, and supernumerary centrosome de-clustering as the possible causes of spindle multipolarity in Myo10 KO HeLa cells.

To define the causes of spindle multipolarity in Myo10 KO HeLa cells, we used staining for γ-tubulin and the centriole marker centrin-1 (along with DAPI), as this staining reveals MTOCs arising from both acentriolar PCM fragments and centriole-containing structures. Imaging of unsynchronized, metaphase Myo10 KO HeLa cells stained in this way revealed examples of multipolar spindles caused by all three mechanisms discussed above. For example, Z-Stack Movie 11, along with the enlarged images of this cell’s three presumptive spindle poles (Fig. 6A1), represents an example of spindle multipolarity driven by centriole disengagement (note that two of this cell’s three presumptive spindle poles possess only one centriole). Z-Stack Movie 12, along with the enlarged images of this cell’s four presumptive spindle poles (Fig. 6A2), represents an example of spindle multipolarity driven by PCM fragmentation (note that two of this cell’s four presumptive spindle poles are γ-tubulin-positive foci that lack centrioles). More extreme examples of multipolar spindles driven by PCM fragmentation were also seen (Fig. S3A and Z-Stack Movie 18; Fig. S3B and Z-Stack Movie 19). Finally, the still images in Figure 6, Panel A3, show an example of spindle multipolarity driven by an inability to cluster supernumerary centrosomes (i.e. centrosome de-clustering; note that this cell contains four presumptive spindle poles, each possessing two centrioles). Examples of partial supernumerary centrosome de-clustering were also observed (Fig. 6A4; note that only two of this cell’s three presumptive spindle poles (the two with four centrioles) were created by supernumerary centrosome clustering).

**Figure 6.**
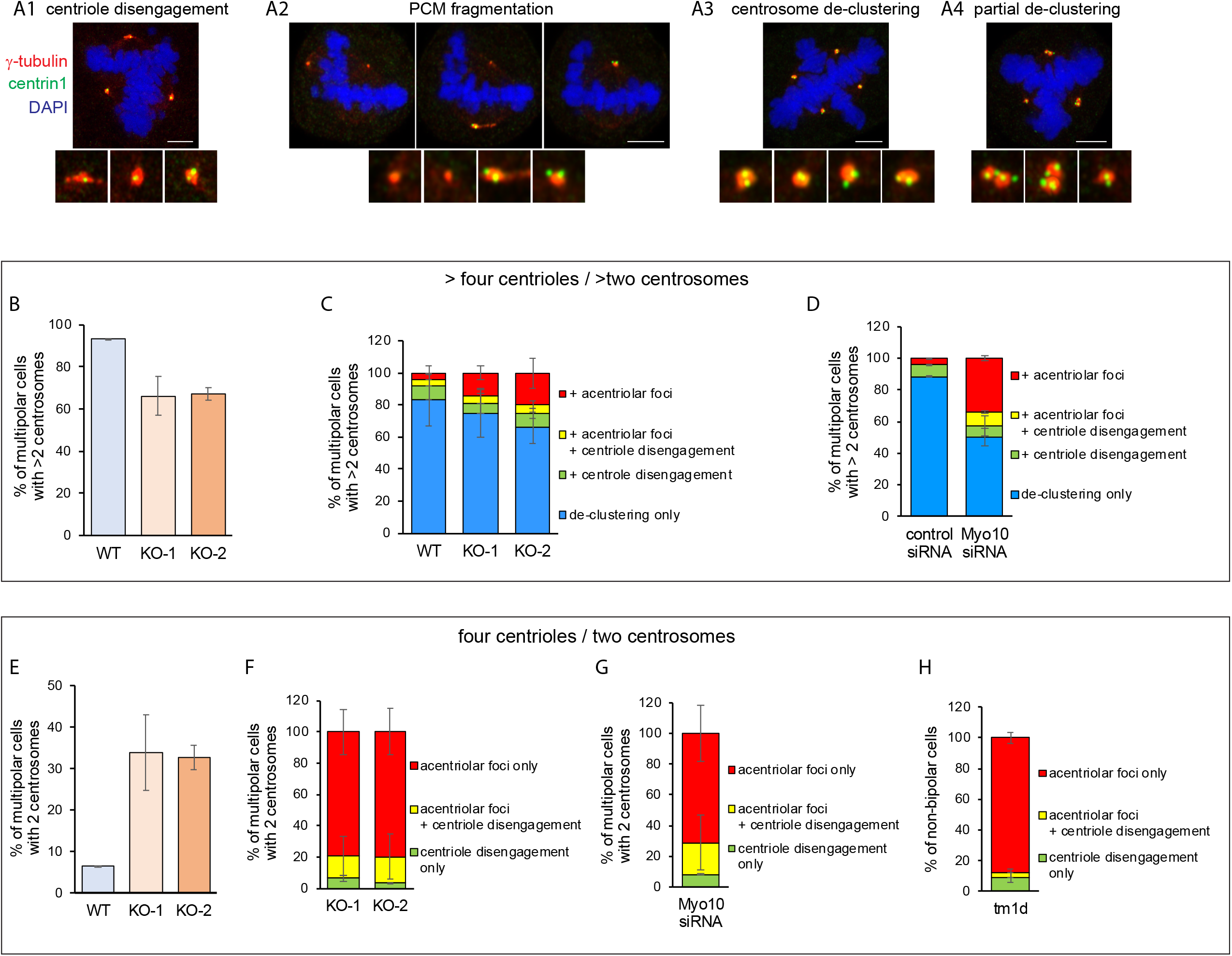
Causes of spindle multipolarity in Myo10-depleted HeLa cells and Myo10 KO MEFs. (A1-A4). Representative images of cells stained for g-tubulin, centrin-1 and DNA that exhibit centriole disengagement (A1), PCM fragmentation (A2; shown are three different focal planes in one cell), centrosome de-clustering (A3), and partial centrosome de-clustering (A4). (B) Percent of multipolar WT HeLa, KO-1 and KO-2 that have more than 4 centrioles/2 centrosomes. (C) Percent of multipolar WT HeLa, KO-1 and KO-2 with more than 4 centrioles/2 centrosomes that exhibit centrosome de-clustering only (blue), centrosome de-clustering plus acentriolar foci (red), centrosome de-clustering plus centriole disengagement (green), or centrosome de-clustering plus acentriolar foci and centriole disengagement (yellow). (D) Percent of multipolar HeLa cells exhibiting the mitotic defects described in Panel C that had been treated with control siRNA or Myo10 siRNA and that had more than 4 centrioles/2 centrosomes. (E) Percent of multipolar KO-1 and KO-2 with 4 centrioles/2 centrosomes. (F) Percent of multipolar WT HeLa, KO-1 and KO-2 with 4 centrioles/2 centrosomes that exhibit acentriolar foci (red), acentriolar foci plus centriole disengagement (yellow), or centriole disengagement only (green). (G) Percent of multipolar HeLa cells exhibiting the mitotic defects described in Panel F that had been treated with Myo10 siRNA-treated and that had 4 centrioles/2 centrosomes (H) Percent of non-bipolar (semipolar plus multipolar) Myo10 KO MEFs isolated from the tm1d mouse that exhibit the mitotic defects described in Panel F. All mag bars are 5 μm.

To quantitate the relative contributions made by centriole disengagement, PCM fragmentation, and supernumerary centrosome de-clustering to the multipolar spindle phenotype, we divided WT and Myo10 KO HeLa cells exhibiting multipolar spindles into two groups: those with more than four centrioles (i.e. cells that possessed supernumerary centrosomes) and those with four centrioles (i.e. cells that did not possess supernumerary centrosomes). For WT Hela, the vast majority (93.4 ± 0.1%) contained more than four centrioles (Fig. 6B). In 82.9 ± 16.3% of these cells, every γ-tubulin spot contained at least two centrioles (Fig. 6C; WT blue), indicating that centrosome de-clustering was solely responsible for their multipolar spindle phenotype. For the remaining 17.1% of multipolar WT HeLa with more than four centrioles, 8.5 ± 8.0% exhibited centrosome de-clustering plus centriole disengagement (one or more γ-tubulin spots containing only one centriole) (Fig. 6C; WT green), 4.3 ± 4.0% exhibited centrosome de-clustering plus acentriolar foci arising (one or more γ-tubulin spots containing no centrioles) (Fig. 6C; WT red), and 4.3 ± 4.0% exhibited centrosome de-clustering plus both centriole disengagement and acentriolar foci (Fig. 6C; WT yellow).

For KO-1 and KO-2 cells, about two thirds contained more than four centrioles (66.1 ± 9.2% for KO-1 and 67.3 ± 3.0% for KO-2) (Fig. 6B). In roughly two thirds of these cells, every γ-tubulin spot contained at least two centrioles (74.9 ± 15.0% for KO-1 and 65.9 ± 9.9% for KO-2) (Fig. 6C, KO-1 blue and KO-2 blue), indicating that centrosome de-clustering was mainly responsible for their multipolar spindle phenotype. Like WT HeLa, the remaining one third of multipolar KO cells with more than four centrioles exhibited centrosome de-clustering plus either centriole disengagement, acentriolar foci, or both (Fig. 6C; green, red and yellow, respectively, for KO-1 and KO-2), although the percent of KO-1 cells and KO-2 cells exhibiting acentriolar foci was 3.4-fold and 4.7-fold higher than in WT HeLa cells, respectively (Fig. 6C; compare the red in KO-1 and KO-2 to the red in WT). These results were supported by scoring those WT HeLa cells treated with Myo10 siRNA that contained more than four centrioles, which exhibited a 8.7-fold increase in the percentage of cells with acentriolar foci compared to HeLa cells treated with a control siRNA (Fig. 6D; compare the red in Myo10 siRNA to the red in control siRNA).

Finally, and most interestingly, were the results for cells exhibiting multipolar spindles that contained only four centrioles (i.e. cells that did not possess supernumerary centrosomes), which corresponded to about one third of Myo10 KO cells (33.9 ± 9.2% for KO-1 and 32.7 ± 3.0% for KO-2) but only a tiny fraction of WT HeLa cells (6.6 ± 0.1%) (Fig. 6E). Importantly, acentriolar foci arising from PCM fragmentation was by itself responsible for about ∼80% of the multipolar phenotype exhibited by this group of KO cells (79.4 ± 14.3% for KO-1 and 79.6 ± 14.7% for KO-2) (Fig. 6F; red in KO-1 and KO-2). Of the remaining ∼20% of multipolar KO cells with four centrioles, about three quarters exhibited acentriolar foci along with centriole disengagement (Fig. 6F; yellow in KO-1 and KO-2). In total, therefore 93.6% of multipolar KO-1 cells lacking supernumerary centrosomes, and 96.4% of KO-2 cells lacking supernumerary centrosomes, exhibited acentriolar foci, arguing that PCM fragmentation is the primary cause of spindle multipolarity in these cells. Importantly, very similar results were obtained for HeLa cells subjected to transient Myo10 depletion using Myo10 siRNA. Specifically, 91.9% of multipolar Myo10 KD cells that did not possess supernumerary centrosomes exhibited acentriolar foci arising from PCM fragmentation (Fig. 6G; red plus yellow in Myo10 siRNA). Together, these results argue that Myo10 supports spindle bipolarity in HeLa cells by promoting both PCM integrity and the clustering of supernumerary centrosomes.

### Multipolar spindles in Myo10 KO MEFs arise primarily from PCM fragmentation

Staining of MEFs isolated from the straight Myo10 KO mouse (tm1d) that had only four centrioles/two centrosomes showed that acentriolar foci arising from PCM fragmentation was by itself responsible for 88.0% of the multipolar phenotypes exhibited by this group of cells (Fig. 6H; red). Of the remaining 12.0% of multipolar KO MEFs with four centrioles, about one third exhibited acentriolar foci plus centriole disengagement (Fig. 6H; yellow). In total, therefore, ∼91% of multipolar KO MEFs lacking supernumerary centrosomes exhibited acentriolar foci, arguing that PCM fragmentation is the primary cause of spindle multipolarity in these cells.

### Acentriolar foci arising from PCM fragmentation serve as MTOCs during mitosis

The preceding analyses of multipolar cells were done using unsynchronized cells stained for DNA (DAPI), centrioles (centrin-1) and γ-tubulin. Presumptive spindle poles were identified in these images based on DNA organization and on staining for γ-tubulin. In addition, over-exposure of the γ-tubulin signal often revealed evidence of microtubule asters emanating from these presumptive spindle poles. To provide additional evidence that PCM fragments lacking centrioles serve as spindle poles, we transfected Myo10 KO HeLa cells with dTomato-centrin-1 to mark centrioles, H2B-iRFP670 to mark chromatin, and EB1-EGFP to identify all MTOCs (i.e. those with centrioles and those lacking centrioles because they were created by PCM fragmentation). As expected, Myo10 KO HeLa cells containing two centriolar poles and undergoing normal bipolar mitoses exhibited EB1-EGFP comets emanating from two centriolar spindle poles (Fig. 7A1, A2 and Z-Stack Movie 13). Also as expected, Myo10 KO HeLa cells containing three centriolar poles and undergoing multipolar mitoses exhibited EBI-EGFP comets emanating from three centriolar spindle poles (Fig. 7B1, B2 and Z-Stack Movie 14). Importantly, imaging EBI-EGFP in Myo10 KO HeLa undergoing multipolar mitoses also revealed microtubule asters emanating from acentriolar poles in addition to centriolar poles (Fig. 7C1, C2 and Z-Stack Movie 15; enlarged images of centriolar and acentriolar spindle poles are shown in Movies 16 and 17, respectively). Together, these results confirm that PCM fragmentation creates acentriolar spindle poles that contribute to the multipolar phenotype exhibited by cells lacking Myo10.

**Figure 7.**
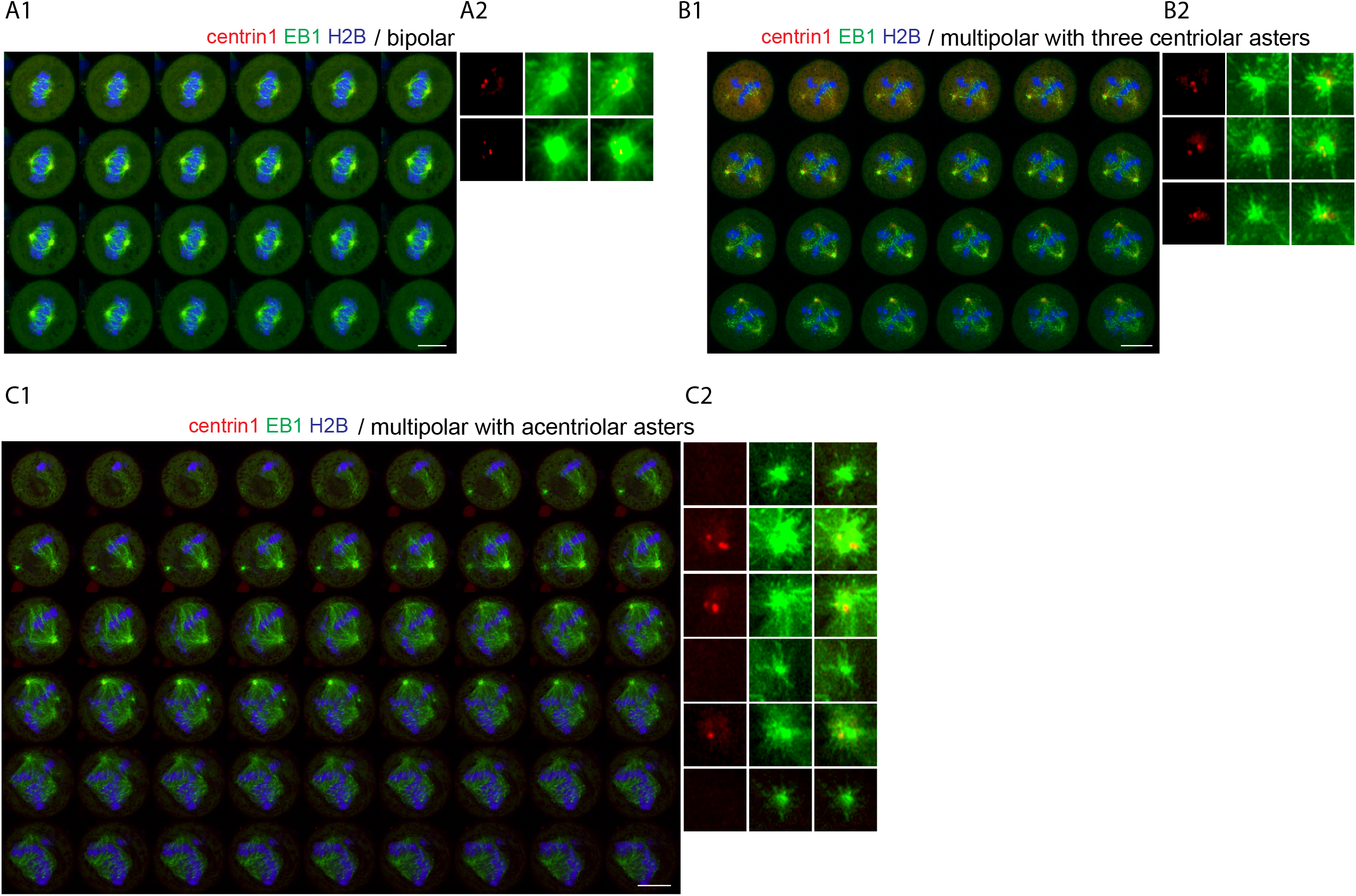
Acentriolar foci arising from PCM fragmentation serve as MTOCs during mitosis. (A1 and A2) Optical sections (A1) of a KO-1 cell containing two centriolar poles and undergoing a normal bipolar mitosis that had been transfected with EB1-EGFP to identify all MTOCs, dTomato centrin-1 to mark centrioles, H2B-iRFP670 to mark chromatin (see also Z-Stack Movie 13). The insets (A2) show enlargements of this cell’s two poles. (B1 and B2) Same has A1 and A2 except that this KO-1 cell contained three centriolar poles and was undergoing a multipolar mitosis (see also Z-Stack Movie 14). (C1 and C2) Same as A1 and A2 except that this KO-1 cell contained three acentriolar poles (i.e. negative for dTomato centrin-1) generated by PCM fragmentation in addition to three centriolar poles (see also Z-Stack Movie 15). Movies 16 and 17 show examples of EB1-EGFP comets emanating from a centriolar pole and an acentriolar pole, respectively. All the mag bars are 10 μm.

### PCM/pole fragmentation occurs primarily at metaphase, arguing that it is a force-dependent event

Defects in spindle pole integrity that lead to PCM/pole fragmentation are commonly revealed around metaphase when the chromosomal and spindle forces that the pole must resist increase [47]. To determine when the PCM fragments in Myo10 KO cells, we stained unsynchronized, mitotic KO cells for γ-tubulin, centrin-1 and DAPI. Scoring the percent of cells containing only two centrosomes that exhibited y-tubulin-positive, centriole-negative PCM fragments showed that these fragments only begin to appear at prometaphase and peak in frequency at metaphase (Fig. 8A). These results indicate that cells do not enter mitosis with acentriolar PCM fragments, and that PCM/pole fragmentation occurs primarily between prometaphase and metaphase, consistent with it being a force-dependent event.

**Figure 8.**
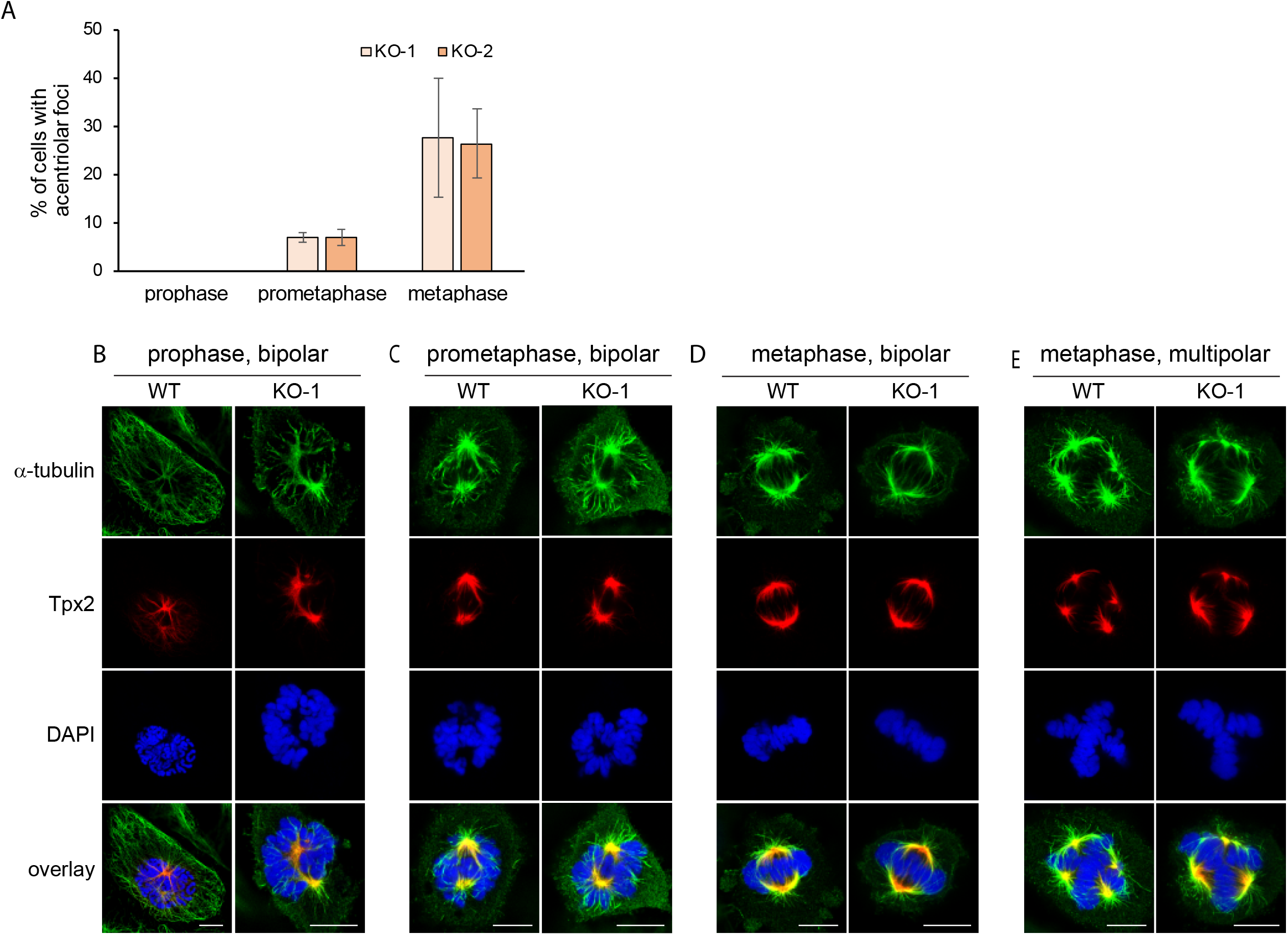
PCM/pole fragmentation is a force-dependent event that is not due to a defect in TPX2 localization. (A) Shown is the percent of unsynchronized KO-1 and KO-2 cells containing only two centrosomes that exhibited y-tubulin-positive, centriole-negative PCM fragments at prophase, prometaphase and metaphase. (B-E) Representative images of WT HeLa cells and Myo10 KO HeLa cells at prophase (B), prometaphase (C), and metaphase (D, E)) that were stained for α-tubulin, TPX2 and DNA (plus overlay) while undergoing bipolar divisions (B-D) or a multipolar division (E). All the mag bars are 5 μm.

### PCM fragmentation in Myo10 KO HeLa cells is not due to a defect in the pole localization of TPX2

As reviewed in the Introduction, Bement and colleagues [42] showed previously that embryonic frog epithelial cells depleted of Myo10 using morpholinos exhibit spindle pole fragmentation at metaphase, leading to an increase in the frequency of multipolar spindles. Moreover, they attributed this apparent defect in pole integrity to a defect in the localization of the spindle pole assembly factor TPX2 at and near poles when Myo10 is depleted. To determine whether the PCM fragmentation observed here is also caused by a defect in TPX2 localization, we stained WT and Myo10 KO HeLa cells for α-tubulin, TPX2 and DNA at prophase, prometaphase and metaphase. Figure 8, Panels B-D, show that KO cells undergoing bipolar mitosis do not exhibit any obvious defect in the localization of TPX2 at and near spindle poles at all three mitotic stages. Similarly, the localizations of TPX2 in WT and KO HeLa cells undergoing multipolar mitosis are indistinguishable (Fig. 8E). These results, together with that fact that Myo10 is rarely seen at poles in HeLa cells expressing mScarlet-Myo10 (Fig. 4), argues that the PCM fragmentation observed here cannot be attributed to Myo10 acting alone at poles or through TPX2 recruitment to maintain pole integrity (although see Discussion).

### The defect in PCM/pole integrity in Myo10 KO HeLa cells is not due to an obvious defect in spindle pole maturation

Defects in spindle pole maturation can lead to defects in pole integrity [47]. To look for a possible defect in spindle pole maturation in Myo10 depleted cells, we stained WT, KO-1 and KO-2 HeLa cells at prophase, prometaphase and metaphase for centrin-1, DNA, and the pole protein CDK5Rap2, which is known to accrue in maturing poles where it recruits γ-TuRC to promote microtubule nucleation [45, 46]. Stained cells possessing only two poles were optically sectioned in 0.25 μm intervals and the total intensity of the CDK5Rap2 signal obtained by summing slices (see Methods for additional details). Representative images of stained poles in KO-1 cells at prophase, prometaphase and metaphase are shown on Figure 9, Panels A-C, respectively. Importantly, quantitation revealed no significant differences between WT cells and either KO-1 or KO-2 in the pole content of CDK5Rap2 at all three mitotic phases (Fig. 9D). This result argues that Myo10’s contribution to spindle pole integrity may involve a role in maintaining pole stability rather than promoting pole maturation (see Discussion).

**Figure 9.**
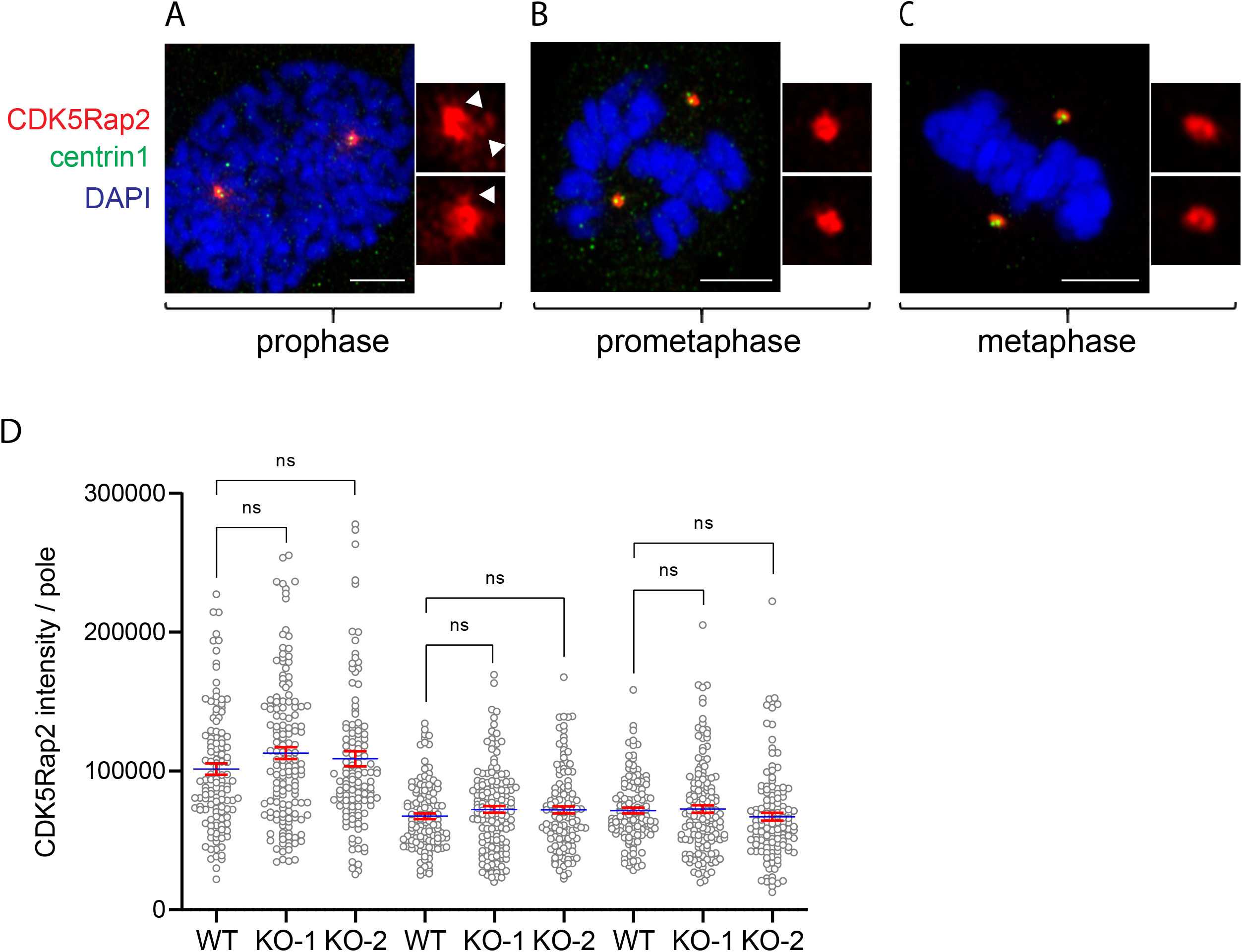
The defect in PCM/pole integrity in Myo10 knockout cells is not due to an obvious defect in spindle pole maturation. (A-C) Representative images of unsynchronized KO-1 cells stained at prophase (A), prometaphase (B), and metaphase (C) for centrin-1, DNA, and the pole protein CDK5Rap2. The enlarged insets show the signal for CDK5Rap2 only (the white arrowheads in the insets for (A) point to CDK5Rap2-postive pericentriolar satellites). (D) Quantitation of the amount of CDK5Rap2 at poles in WT HeLa, KO-1 and KO-2 at prophase, prometaphase and metaphase. The fact that the pole content of CDK5Rap2 is higher at prophase than at prometaphase and metaphase may be due to the presence of CDK5Rap2-positive pericentriolar satellites surrounding the poles at prophase but not at prometaphase and metaphase. All mag bars are 5μm.

## Discussion

Here we showed that the primary driver of spindle multipolarity exhibited by Myo10 KO HeLa cells lacking supernumerary centrosomes and by Myo10 KO MEFs is PCM fragmentation, which creates y-tubulin-positive, centriole-negative microtubule asters that serve as additional spindle poles. We also showed that Myo10 depletion further accentuates spindle multipolarity in HeLa cells possessing supernumerary centrosomes by impairing centrosome clustering. Importantly, all of our scoring was done using unsynchronized cells, as treatments used to synchronize cells by delaying mitotic entry (e.g. low dose nocodazole) can lead to the formation of multipolar spindles with abnormal centriole distributions at poles (e.g. single or no centrioles) due to cohesion fatigue [47].

PCM fragmentation is one of four ways to get multipolar spindles in the absence of centrosome amplification (reviewed in [47]). Importantly, we showed that two of the other three ways-cytokinesis failure and centriole overduplication-did not contribute to the multipolar phenotype exhibited by Myo10 KO cells, and that the remaining way-centriole disengagement-played only a minor role. PCM fragmentation typically occurs when the physical integrity of spindle poles is compromised in some way, causing them to fragment when subjected to the strong pulling forces that develop during mitosis, and resulting in the accumulation of one or more acentriolar poles in addition to two centriole-containing poles. While there are numerous examples of pole defects that give rise to PCM fragmentation (reviewed in [47]), the study that is most relevant to our work is that of Bement and colleagues [42], who showed that embryonic frog epithelial cells depleted of Myo10 enter mitosis as bipolar but become multipolar during anaphase due to pole fragmentation (of note, they did not stain for centrioles, so it is unclear to what extent pole fragmentation in their cells was due to centriole disengagement versus the generation of acentriolar poles by PCM fragmentation). Bement and colleagues attributed the apparent defect in spindle pole integrity to a defect in the recruitment of the spindle pole assembly factor TPX2, which they argued is recruited to poles through an interaction with Myo10’s MyTH4/FERM domain. We, on the other hand, did not see any obvious reduction in the localization of endogenous TPX2 at and near spindle poles in our Myo10 KO HeLa cells. This result, together with the fact that Myo10 was rarely seen at spindle poles in HeLa cells, argues that the PCM fragmentation observed here cannot be attributed to Myo10 acting alone at poles or through TPX2 recruitment to maintain pole integrity. That said, we cannot completely exclude the possibility that a small and/or transient pool of Myo10 at spindle poles serves to maintain spindle pole integrity, or that we missed subtle but functionally important differences in the pole localization of TPX2 in Myo10 KO cells.

We showed that PCM fragmentation in Myo10 KO HeLa cells entering mitosis as bipolar occurs primarily between prometaphase and metaphase, which is when the chromosomal and spindle forces placed on spindle poles increase. This observation is consistent with the force-dependent fragmentation of poles whose physical integrity is somehow compromised when Myo10 is missing. One problem that can lead to defects in pole integrity is a defect in pole maturation [45-47]. Quantitation of the content of CDK5Rap2 at the poles of WT, KO-1 and KO-2 HeLa cells during prophase, prometaphase and metaphase did not, however, reveal an obvious defect in pole maturation when Myo10 is missing. That pole maturation, as assessed using a standard marker for this event [45, 46], appears normal in the absence of Myo10 suggests that the myosin might be required instead for the stability of poles. In *C. elegans* embryos, poles are thought to undergo a major transition in their physical properties wherein they go from a strong, ductile state in metaphase to a weak, brittle state in anaphase [64, 65]. These two states underlie two of the pole’s essential functions during mitosis: the ability to resist strong, microtubule-dependent pulling forces during metaphase, and the ability to undergo microtubule force-dependent PCM rupture, a central feature of the pole disassembly process that begins at anaphase. This transition is thought to be regulated by the localization of activated PLK-1 and SPD-2 to poles, wherein phosphorylated PLK-1 and SPD-2 present in metaphase poles confer strength and ductility to the PCM scaffold, and their PP2A phosphatase-mediated dissociation from poles at anaphase causes the poles to transition to a weak, brittle state that permits force-dependent PCM fragmentation and pole disassembly [64, 65]. One possibility, then, is that Myo10 helps in some way to reinforce the PCM scaffold such that in its absence, spindle poles would transition prematurely to their weak, brittle state. If this were to occur during metaphase, the ensuing destabilization of the poles could lead to PCM fragmentation and a multipolar spindle phenotype. Since Myo10 does not localize robustly to spindle poles, we speculate that it would act indirectly through other PCM-resident regulators or structural components. Future efforts should seek to determine if Myo10 does in fact influence the time when poles transition from strong and ductile to a weak and brittle, and if so, how.

Another explanation for how Myo10 might promote PCM/pole integrity without localizing to poles involves its ability regulate the CDK1 kinase inhibitor WEE1 [60]. Normally, CDK1 activity increases during prophase, is highest at metaphase, and then decreases at anaphase onset to promote mitotic progression [66]. Importantly, Bement and colleagues have presented evidence that Myo10 binds WEE1 as cells approach metaphase to prevent WEE1 from inactivating CDK1, and then releases WEE1 at the end of metaphase so that it can suppress CDK1 activity and promote mitotic progression [60]. In this way, then, the interaction of Myo10 with WEE1 would coordinate Myo10’s role in attaining correct spindle positioning and orientation [42] with anaphase onset. One interesting ramification of this model has to do with the role that CDK1 activity plays in determining the localization of the microtubule-binding and dynein-anchoring protein NuMA [66-69]. Normally, the robust activity of CDK1 up to metaphase keeps NuMA largely phosphorylated, which in turn keeps it largely localized to spindle poles, where is promotes pole focusing and pole stability, and in the spindle, where in promotes load sharing by crosslinking k-fibers to interpolar microtubules (although some unphosphorylated NuMA present in the cortex serves to promote metaphase spindle positioning by recruiting dynein) [66-70]. At anaphase onset, however, CDK1 activity drops. This drop, together with the action of the phosphatase PPP2CA, causes NuMA’s phosphorylation state to drop. This precipitates the translocation of significant amounts of NuMA from spindle poles and the spindle to the cortex, where it, together with bound dynein, drives spindle elongation during anaphase [66-70]. Given all this, one would predict that in the absence of Myo10-dependent WEE1 sequestration (i.e. in Myo10 KO cells), CDK1 would be inhibited prematurely, leading to the premature translocation of NuMA from spindle poles and the spindle to the cortex. This could in turn lead to premature changes in pole stability and spindle load sharing that, together with premature escalation of cortical pulling forces, could lead to PCM/pole fragmentation at metaphase. Future efforts should examine NuMA’s dynamic localization across mitosis in cells lacking Myo10, and define the physiological significance of Myo10-WEE1 interaction by complementing Myo10 KO cells with a version of Myo10 that cannot bind WEE1.

While our data indicated that PCM fragmentation contributes significantly to the formation of multipolar spindles in Myo10 KO HeLa cells possessing supernumerary centrosomes, it indicated that the primary cause of spindle multipolarity in these cells is an inability to cluster their extra centrosomes. This result confirms and extends the seminal observations made by Kwon and colleagues using RNAi screens for genes whose expression promotes supernumerary centrosome clustering, which implicated Myo10 in MDA-231 cancer cells and fly Myo10A (a homolog of the human MyTH4/FERM myosin, Myo15) in near-tetraploid *Drosophila* S2 cells [44]. Importantly, this screen also identified the pole focusing, microtubule minus end-directed kinesin 14 family member HSET (Ncd in Drosophila) as being essential for supernumerary centrosome clustering. While it seems obvious that a pole-focusing microtubule motor would be required to cluster extra poles, how Myo10 cooperates with HSET to promote clustering is not as obvious. One possibility is that Myo10 uses its ability to engage microtubules via it’s microtubule-binding MyTH4 domain, together with its ability as a myosin to localize to the actin-rich cortex, to facilitate supernumerary centrosome clustering. Notably, this mechanism would be similar to the mechanism proposed by Kwon and colleagues [43] for how Myo10 cooperates with dynein to position the metaphase spindle.

We presented data here that, together with other published data, supports an adhesion-centric mechanism for how Myo10 might promote supernumerary centrosome clustering. First, we showed that Myo10 localizes robustly to retraction fibers in mitotic HeLa cells. These actin-based structures have been considered for decades as being essential for the adhesion of cells dividing in 2D [54-58, 71]. Second, we showed that the spindle in Myo10 KO cells is often not parallel to the substratum. This result, and a similar result reported by Toyoshima and colleagues in Myo10 knockdown NRK cells [51], is consistent with Myo10 depletion causing a defect in cell adhesion during mitosis. Importantly, this conclusion is completely in line with the now strong evidence that Myo10 supports the formation of adhesions within filopodia and lamellipodia in interphase cells by virtue of its FERM domain-dependent interaction with β1 integrin [12, 23-25]. Moreover, it is in line with micropatterning data showing that the ability of cancer cells to cluster their supernumerary centrosomes is influenced significantly by the pattern of adhesion [44]. Together, these results point towards an adhesion-based mechanism wherein Myo10’s ability to promote adhesion during mitosis promotes supernumerary centrosome clustering by anchoring and possibly orienting the pole-focusing forces exerted by HSET. Given that retraction fiber-based adhesions are known to play a central role in positioning the mitotic spindle [72-74], it may also be the case that Myo10 promotes spindle positioning at least in part by promoting cell adhesion during mitosis.

If not clustered, supernumerary centrosomes cause multipolar cell divisions that result in chromosome miss-segregation, aneuploidy and cell death [61-63]. Given this, and given that supernumerary centrosomes are rare in normal cells and common in cancer cells, drugs that inhibit the clustering of supernumerary centrosomes should selectively kill cancer cells. Indeed, compounds that inhibit HSET are currently being tested as anticancer therapeutics [75]. By the same token, a specific inhibitor of Myo10 might also serve to selectively kill cancer cells. Moreover, it might work synergistically with inhibitors of HSET. This seems likely given the results of Kwon et al [44], who showed using assays that quantified the suppression of supernumerary centrosome clustering that actin disassembly was not synergistic with Myo10 knockdown (presumably because they act in the same pathway), but was synergistic with HSET inhibition. Notably, several recent studies have provided additional support for the idea that inhibitors of Myo10 might be effective as cancer therapeutics. First, Collucio and colleagues [76] reported that depletion of Myo10 in mouse models of melanoma reduces melanoma development and metastasis and extends medial survival time. Similarly, Rosenfeld and colleagues [77] reported that the lifespan of mice with glioblastoma is extended significantly on a Myo10 KO background. Third, Zhang and colleagues [78] reported that the progression of breast cancer tumors in mice is inhibited by Myo10 depletion and accelerated by Myo10 over expression. More generally, many cancer cell types exhibit elevated levels of Myo10, which may help support their growth by promoting the clustering of their extra centrosomes [40, 79, 80]. These and other studies [40], together with the data presented here, provide strong justification for performing screens to identify inhibitors of Myo10 as possible cancer therapeutics.

Finally, while studies performed over the last several decades using cells dividing in 2D have provided a wealth of mechanistic information, mitosis in 3D (i.e. in tissue) presents the more physiological context in which to define such mechanisms (reviewed in [81-85]). Importantly, cells dividing in 3D have constraints on their shape and additional spatial cues provided by intercellular adhesions that cells in 2D lack. Along these lines, we found that Myo10 KO MEFs proliferate faster when maintained at high density, suggesting that spatial cues provided by neighboring cells even in 2D may might mitigate to some extent the defects in mitosis caused by the loss of Myo10. That said, Cheney and colleagues reported that MDCK cells in monolayer exhibit significant defects in spindle orientation when Myo10 is knocked down [86]. Similarly, Bement and colleagues showed that cells within embryonic frog epithelia exhibit numerous mitotic defects when Myo10 is depleted [42]. These studies argue, then, that Myo10 makes significant contributions to mitosis even in the context of a 3D tissue environment. To further clarify Myo10’s contribution to mitosis in this context, our future efforts will include characterizing mitosis in intestinal organoids prepared from Myo10 KO mice.

## Materials and Methods

### Cell culture and transfection

HeLa cells (ATCC, CCL-2), HEK-293T cells (ATCC, CRL-3216), and primary MEFs were cultured in high glucose DMEM (Gibco) supplemented with 10% FBS (Gibco) and Antibiotic-Antimycotic solution (Gibco), and were maintained at 37°C in a 5% CO_2_ humidified incubator. MEFs were prepared from ∼E19 embryos as described previously [87]. Only low passage number MEFs (<P3) were used for quantitating knockout phenotypes. For imaging purposes, cells were cultured in the coverglass bottom chamber slides (Cellvis) coated with Fibronectin (10 ng/ml; Gibco). Cells were transfected using either Lipofectamine 3000 (Invitrogen) or an Amaxa nucleofection apparatus (Lonza) according to the manufacturer’s instructions. To knock down Myo10, the ON-TARGETplus human Myo10 siRNA-SMARTpool from Dharmacon RNA Technologies (L-007217-00) was transfected into WT Hela cells using Lipofectamine RNAiMAX reagent (Invitrogen). To increase knockdown efficiency, the transfection was carried out twice with two-days interval in a reverse way.

### Mice

WT C57BL/6 mice was purchased from Jackson Laboratories (#000664). The creation of the straight Myo10 KO mouse (tm1d) and the Myo10 cKO mouse (tm1c) is described in [13]. Both Myo10 KO mice are on a pure B6 background. All animal experiments were approved by the Animal Care and Use Committee of the National Heart, Lung, and Blood Institute, in accordance with the National Institutes of Health guidelines.

### Antibodies and immunofluorescence

The following primary antibodies were used: mouse anti-γ-tubulin (MilliporeSigma, T6793, 1:300), mouse anti-α-tubulin (Abcam, ab7291, 1:300), rabbit anti-α-tubulin (Abcam,ab52866, 1:300), rabbit anti-myo10 (MilliporeSigma, HPA024223, 1:300), mouse anti-centrin1 (MilliporeSigma, 04-1624, 1:200), rabbit anti-centrin-1 (Abcam, ab101332, 1:200), rabbit anti-CDK5Rap2 (MilliporeSigma, 06-1398, 1:200), rabbit anti-pericentrin (Abcam, ab4448, 1:200), mouse anti-β-actin (Abcam, ab6276, 1:10,000), and mouse anti-Tpx2 (Santa Cruz Biotechnology, sc-53775, 1:200). AlexaFluor-conjugated and HRP-conjugated secondary antibodies were purchased from Jackson ImmunoResearch Laboratories. DAPI, Alexa Fluor 488 labeled Phalloidin, and Alexa Fluor 568 labeled Phalloidin were purchased from ThermoFisher Scientific. Cells were fixed in 4% PFA for 10 min at RT unless subsequent staining was for centrosomes and pole-related proteins, in which case the cells were subjected to a two-step fixation method involving 1.5% PFA for 5 min followed by ice-cold MeOH for 5 min at -20°C. All PFA solutions were prepared in Cytoskeleton Stabilization Buffer (150 mM NaCl, 5 mM EGTA, 5 mM glucose, 5 mM MgCl_2_, and 10 mM PIPES (pH 6.8)). Fixed cells were permeabilized and blocked by incubation for 15 min in PBS containing 0.15% Saponin and 5% fetal bovine serum (FBS). Fixed, permeabilized cells were incubated at RT for 1 hr with primary and secondary antibodies diluted in blocking solution, with an intervening wash cycle using PBS.

### Lentivirus packaging and cell transduction

We used the second generation lentiviral packaging plasmid psPAX2 (Addgene, #12260) to generate viral supernatants for pLenti-EB1-EGFP (Addgene, #118084), pLVX-FLAG-dTomato-centrin-1 (Addgene,#73332), pLentiPGK DEST H2B-iRFP670 (Addgene, 90237), and pLenti-EGFP-Cre (Addgene, 86805). These plasmids were cotransfected with psPAX2 and pMD2.G (Addgene, #12259) into HEK-293T cells at ∼80% confluency using LipoD293 (SignaGen, SL100668). Viral supernatants were collected 48 hrs post transfection, clarified by centrifugation at 500 x g for 10 min, and concentrated using a Lenti-X concentrator (TakaraBio, #631231). Viral pellets were obtained by centrifugation of the concentrated virus at 1,500 x g for 45 minutes at 4°C. The pellets were resuspended in complete DMEM and stored in aliquots at -80°C. Purified lentivirus was added directly to cells after the optimal amount was determined by a pilot experiment involving serial dilutions and determining the fraction of cells transduced. After a 6 hrs incubation, the cells were washed 3 times with PBS and returned to complete culture medium for imaging or fixation at various time points. To knock out Myo10 in MEFs isolated from the Myo10 cKO mouse [13], the MEFs were treated with lentivirus expressing EGFP-Cre as described above. Cell phenotypes were determined 48 hrs after lenti-cre transduction. All processes and materials were handled in accordance with the NIH biosafety guidelines.

### Generation of Myo10 KO HeLa cells

To knock out Myo10 in HeLa cells using CRISPR, we used an online CRISPR design tool (http://cripr.mit.edu) to identify the following guide sequences within exon 3 of human Myo10 (NCBI Reference Sequence: NC_000005.10): 5’-CACCGTATGCACCCCACGAACGAGG (PAM)-3’; 3’-CATACGTGGGGTGCTTGCTCCCAAA-5’. These guides sequences were inserted into pSpCas9(BB)-2A-GFP (Addgene, #48138) to create pSpCas9(BB)-2A-GFP-ghMyo10 as described previously [88].Two days after transfection of WT HeLa cells with pSpCas9(BB)-2A-GFP-ghMyo10 using Amaxa nucleofection (Lonza), GFP-positive cells were subjected to single-cell sorting into 96-well plates using a BD FACS cell sorter. Whole cell protein lysates and genomic DNA prepared from the two single-cell KO clones obtained (KO-1 and KO-2) were characterized by Western blotting and sequencing of PCR products that span the guide sequence sites, respectively.

### Immunoblotting

Whole cell protein lysates were collected directly from culture dishes by adding 1X SDS sample buffer, as described previously [89]. Samples were resolved on 4-12% or 6% NuPAGE Bis-Tris gels (ThermoFisher Scientific) and transferred onto nitrocellulose membranes (Bio-Rad) using a semi-dry transfer system (Bio-Rad). Nitrocellulose membranes were blocked in TBST (10 mM Tris, pH 8.0, 150 mM NaCl, and 0.02% Tween 20) supplemented with 5% milk for 2 hrs, incubated with anti-Myo10 primary antibody overnight at 4°C, washed with TBST, and incubated in secondary antibody at RT for 2 hrs. Antibodies were diluted in TBST containing 5% milk. Actin detected using an anti-β-actin antibody was used as a loading control. Proteins were detected using SuperSignal West Pico Plus Chemiluminescent Substrate (ThermoFisher Scientific) and quantitated using an Amersham Imager 600 (GE Healthcare Life Sciences).

### Growth rate measurements

Primary MEFs isolated from WT C57BL/6 mouse embryos, non-exencephalic Myo10 KO mouse embryos (tm1d), and exencephalic Myo10 KO mouse embryos (tm1d) were seeded in individual wells of a 6-well plate at 2.50 × 10^5^ cells per well for the dense condition, 1.25 × 10^5^ cells per well for the moderate condition, and 2.5 × 10^4^ cells per well for the sparse condition. Cells were counted using Olympus Cell Counter modal R1 after 3 days of culture.

### Scoring mitotic phenotypes

To score mitotic phenotypes in HeLa cells (WT, Myo10 KO, Myo10 KD, and Myo10 KO cells rescued with pScarlet-Myo10) and MEFs (isolated from WT C57BL/6 mouse embryos, Myo10 KO mouse embryos (tm1d), and Myo10 cKO mouse embryos (tm1c, scored before and after treatment with pLenti-EGFP-Cre)) were plated in imaging chambers at moderate density, cultured overnight, fixed, and stained with anti-γ-tubulin, anti-α-tubulin and DAPI. Of note, cells were never synchronized for these studies. Z-stack images were taken in 0.25 μm steps. The criteria used to score metaphase cells as bipolar, semi-bipolar, or multipolar are described in the text. The same images were used to quantify the percent of cells exhibiting misaligned chromosomes and lagging chromosomes. Quantitation of the distance in Z between the two spindle poles in cells undergoing bipolar mitosis was based on the number of confocal slices between the γ-tubulin foci marking the two poles. To distinguish mitotic cells containing two centrosomes from mitotic cells containing supernumerary centrosomes, and to distinguish γ-tubulin-positive poles containing two centrioles from those containing one centriole or no centriole, metaphase cells were stained with anti-γ-tubulin, anti-centrin-1 and DAPI. To gauge pole maturation, cells were fixed and stained with anti-CDK5Rap2, anti-centrin-1 and DAPI. Cells with a normal number of centrosomes were imaged in 0.25 μm steps, and the total intensity of CDK5Rap2 per pole was determined by summing slices, then thresholding using the otsu function in Image J.

### Dynamic imaging of acentriolar spindle poles using GFP-EB1

Myo10 KO HeLa cells transduced with pLenti-EB1-EGFP, pLVX-FLAG-dTomato-centrin-1, and pLentiPGK DEST H2B-iRFP670 were imaged live on an Airyscan 880 microscope equipped with a 60X, 1.4 NA objective (one frame every 2 sec for 2 min).

### Imaging and statistical analyses

Imaging of both fixed and live cells was performed on a Zeiss Airyscan 880 microscope equipped with a 60X, 1.4 NA objective. Images were processed in auto strength mode using ZenBlack software (Version 2.3) and analyzed using ImageJ. Excel or GraphPad Prism were used for statistical analyses and graphing. Statistical significance was determined using unpaired t-test and indicated as follows: * = P<0.05, ** = P<0.01, *** = P<0.001, and **** = P<0.0001.

## Acknowledgments

This work was supported by the Intramural Research Program of the National Heart, Lung, and Blood Institute (1ZIAHL000514-31 to JAH) and by R01GM134531 to REC. The authors thank Dr. Nasser Rusan, Dr. Jadranka Loncarek, and Dr. Jeffrey B. Woodruff for their expert advice, and Dr. Xufeng Wu and the NHLBI Light Microcopy Core for assistance with imaging.

## Figure Legends

**Figure S1. Myo10 protein levels in Myo10 KO HeLa cells, Myo10 KD HeLa cells, and Myo10 cKO MEFs after treatment with lenti-cre**. (A) The amount of Myo10 in KO-1 and KO-2 expressed as a percent of the amount in WT HeLa cells (N= 6; 7.1 ± 5.2% for KO-1 and 13.8 ± 11.5% for KO-2). (B) The amount of Myo10 in WT HeLa cells 12, 24 and 48 hrs after the start of a second round of treatment with Myo10 siRNA, expressed as a percent of the amount in WT HeLa cells treated with a nontargeting siRNA control (N=3; 8.6 ± 2.1% at 12 hr, 3.4 ± 2.1 % at 24 hr, and 0.3 ± 0.1% at 48 hr). (C) The amount of Myo10 in cKO MEFs 6, 12, 24 and 48 hrs after addition of a lentivirus expressing GFP-tagged Cre recombinase, expressed as a percent of the amount in cKO MEFs before the addition of lenti-cre (N=3; 90.3 ± 6.9% at 0 hr, 33.4 ± 8.1% at 6 hr, 24.6 ± 15.5% at 12 hr, 3.9 ± 3.6% at 24 hr, and 3.1 ± 0.7% at 48 hr). In each case, the amount of Myo10 protein was determined by densitometry of Western blots loaded with whole cell extracts and probed with a Myo10 antibody.

**Figure S2. Cytokinesis failure and centriole overduplication do not contribute to the multipolar phenotype exhibited by Myo10 KO HeLa cells**. (A1 and A2) Unsynchronized WT HeLa, KO-1 and KO-2 were stained with Phalloidin and DAPI (A1 shows the staining of KO-1) and the number of nuclei per cell quantified (A2) (315 WT HeLa, 313 KO-1 cells, and 307 KO-2 cells from 3 experiments; 1.1 ± 0.3 nuclei per cell for WT, 1.0 ± 0.2 nuclei per cell for KO-1, and 1.1 ± 0.3 nuclei per cell for KO-2). (B) Unsynchronized WT HeLa, KO-1 and KO-2 were stained with centrin-1 and the number of centrioles per cell quantified (155 WT HeLa, 154 KO-1 cells, and 172 KO-2 cells from 3 experiments; 3.1 ± 1.6 centrioles per cell for WT, 2.9 ± 1.2 centrioles per cell for KO-1, and 2.9 ± 1.1 centrioles per cell for KO-2).

**Figure S3. Extreme examples Myo10 KO HeLa cells possessing multiple, presumptive, centriole-negative spindle poles generated by PCM fragmentation**. (A and B) Two examples of KO-1 cells stained for γ-tubulin and centrin-1 that exhibit many presumptive, centriole-negative spindle poles generated by PCM fragmentation in addition to two (A) or four (B) normal poles containing two centriole each (see also Z-Stack Movie 18 for (A) and Z-Stack Movie 19 for (B)). All mag bars are 1 μm.

## Movie Legends

**Movie 1**. Z-Stack movie of a bipolar cell at metaphase that was stained with γ-tubulin-cy3, α-tubulin-AF488, and DAPI, and sectioned in 0.25 μm intervals. Played at 5 fps. The mag bar is 5 μm.

**Movie 2**. Z-Stack movie of a semi-bipolar cell at metaphase that was stained with γ-tubulin-cy3, α-tubulin-AF488, and DAPI, and sectioned in 0.25 μm intervals. Played at 5 fps. The mag bar is 5 μm.

**Movie 3**. Z-Stack movie of a multipolar cell at metaphase that was stained with γ-tubulin-cy3, α-tubulin-AF488, and DAPI, and sectioned in 0.25 μm intervals. Played at 5 fps. The mag bar is 5 μm.

**Movie 4**. Z-Stack movie of a bipolar Myo10 KO-1 cell at metaphase that was stained with γ-tubulin-cy3, α-tubulin-AF488, and DAPI, and sectioned in 0.25 μm intervals to show that its two poles are in different Z-planes. Played at 5 fps. The mag bar is 5 μm.

**Movie 5**. Z-Stack movie of a bipolar Myo10 KO-1 cell at anaphase that was stained with γ-tubulin-cy3, α-tubulin-AF488, and DAPI, and sectioned in 0.25 μm intervals to show that its two poles are in different Z-planes. Played at 5 fps. The mag bar is 5 μm.

**Movie 6**. Z-Stack movie of a bipolar Myo10 KO-1 cell at telophase that was stained with γ- tubulin-cy3, α-tubulin-AF488, and DAPI, and sectioned in 0.25 μm intervals to show that its two poles are in different Z-planes. Played at 5 fps. The mag bar is 5 μm.

**Movie 7**. Z-Stack movie of a metaphase Myo10 KO-1 cell expressing mScarlet-Myo10 that was fixed and stained with Alexa488-labeled Phalloidin and DAPI, and sectioned in 0.25 μm intervals to show mScarlet-Myo10 localization at the tips of retraction fibers and dorsal filopodia. Played at 5 fps. The mag bar is 5 μm.

**Movie 8**. Z-Stack movie of a metaphase Myo10 KO-1 cell expressing mScarlet-Myo10 that was fixed and stained with Alexa488-labeled Phalloidin and DAPI, and sectioned in 0.25 μm intervals to show mScarlet-Myo10 localization at/near the poles. Played at 5 fps. The mag bar is 5 μm.

**Movie 9**. Z-Stack movie of a metaphase Myo10 KO-1 cell expressing mScarlet-Myo10 that was fixed and stained with g-tubulin-AF488 and DAPI, and sectioned in 0.25 mm intervals to show mScarlet-Myo10 localization at or very near γ-tubulin-positive spindle poles. Played at 5 fps. The mag bar is 5 mm.

**Movie 10**. Z-Stack movie of a metaphase Myo10 KO-1 cell expressing mScarlet-Myo10 that was fixed and stained with γ-tubulin-AF488 and DAPI, and sectioned in 0.25 μm intervals to show mScarlet-Myo10 at or very near γ-tubulin-positive spindle poles and in spindles. Played at 5 fps. The mag bar is 5 μm.

**Movie 11**. Z-Stack movie of a Myo10 KO-1 cell exhibiting a multipolar spindle produced by centriolar disengagement alone that was stained with γ-tubulin-cy3, centrin-1-AF488 and DAPI, and sectioned in 0.25 μm intervals. Played at 5 fps. The mag bar is 5 μm.

**Movie 12**. Z-Stack movie of a Myo10 KO-1 cell exhibiting a multipolar spindle produced by PCM fragmentation alone that was stained with γ-tubulin-cy3, centrin-1-AF488 and DAPI, and sectioned in 0.25 μm intervals. Played at 5 fps. The mag bar is 5 μm.

**Movie 13**. Live Z-Stack movie of a bipolar Myo10 KO-1 cell with two centriolar poles (2 centrioles/pole) that was expressing EB1-EGFP, dTomato-centrin-1 and H2B-iRFP670, and sectioned in 0.25 μm intervals. Played at 5 fps. The mag bar is 5 μm.

**Movie 14**. Live Z-Stack movie of a multipolar Myo10 KO-1 cell with three centriolar poles (2 centrioles/pole) that was expressing EB1-EGFP, dTomato-centrin-1 and H2B-iRFP670, and sectioned in 0.25 μm intervals. Played at 5 fps. The mag bar is 5 μm.

**Movie 15**. Live Z-Stack movie of a multipolar Myo10 KO-1 cell with three centriolar poles (2 centrioles/pole) and three acentriolar poles that was expressing EB1-EGFP, dTomato-centrin-1 and H2B-iRFP670, and sectioned in 0.25 μm intervals. Played at 5 fps. The mag bar is 5 μm.

**Movie 16**. Time-lapse images of EB1-EGFP at a centriolar pole in the Myo10 KO-1 cell shown in Movie 15. Images were taken every 2 sec for 2 min and are played back at 5 fps. The mag bar is 5 μm.

**Movie 17**. Time-lapse images of EB1-EGFP at an acentriolar pole in the Myo10 KO-1 cell shown in Movie 15. Images were taken every 2 sec for 2 min and are played back at 5 fps. The mag bar is 5 μm.

**Movie 18**. Z-Stack movie of a multipolar Myo10 KO-2 cell with acentriolar poles generated by PCM fragmentation alone that was stained with γ-tubulin-cy3, centrin-1-AF488 and DAPI, and sectioned in 0.25 μm intervals. Played at 5 fps. The mag bar is 5 μm.

**Movie 19**. Z-Stack movie of a multipolar Myo10 KO-2 cell with acentriolar poles generated by PCM fragmentation along with centrosome de-clustering that was stained with γ-tubulin-cy3, centrin-1-AF488 and DAPI, and sectioned in 0.25 μm intervals. Played at 5 fps. The mag bar is 5 μm.

## References

1. Houdusse, A. and M.A. Titus, The many roles of myosins in filopodia, microvilli and stereocilia. Curr Biol, 2021. 31(10): p. R586–R602.

2. Kerber, M.L. and R.E. Cheney, Myosin-X: a MyTH-FERM myosin at the tips of filopodia. J Cell Sci, 2011. 124(Pt 22): p. 3733–3741.

3. Sousa, A.D. and R.E. Cheney, Myosin-X: A molecular motor at the cell’s fingertips. Trends Cell Biol, 2005. 15(10): p. 533–539.

4. Tokuo, H., Myosin X. Adv Exp Med Biol, 2020. 1239: p. 391–403.

5. Weck, M.L., N.E. Grega-Larson, and M.J. Tyska, MyTH4-FERM myosins in the assembly and maintenance of actin-based protrusions. Curr Opin Cell Biol, 2017. 44: p. 68–78.

6. Berg, J.S. and R.E. Cheney, Myosin-X is an unconventional myosin that undergoes intrafilopodial motility. Nat Cell Biol, 2002. 4(3): p. 246–250.

7. Berg, J.S., et al., Myosin-X, a novel myosin with pleckstrin homology domains, associates with regions of dynamic actin. J Cell Sci, 2000. 113 Pt 19(19): p. 3439–3451.

8. Pi, X., et al., Sequential roles for myosin-X in BMP6-dependent filopodial extension, migration, and activation of BMP receptors. J Cell Biol, 2007. 179(7): p. 1569–1582.

9. Alieva, N.O., et al., Myosin IIA and formin dependent mechanosensitivity of filopodia adhesion. Nat Commun, 2019. 10(1).

10. Bohil, A.B., B.W. Robertson, and R.E. Cheney, Myosin-X is a molecular motor that functions in filopodia formation. Proc Natl Acad Sci U S A, 2006. 103(33): p. 12411–12416.

11. Hammers, D.W., et al., Filopodia powered by class x myosin promote fusion of mammalian myoblasts. eLife, 2021. 10: p. 1–50.

12. He, K., et al., Myosin X is recruited to nascent focal adhesions at the leading edge and induces multi-cycle filopodial elongation. Sci Rep, 2017. 7(1).

13. Heimsath, E.G., et al., Myosin-X knockout is semi-lethal and demonstrates that myosin-X functions in neural tube closure, pigmentation, hyaloid vasculature regression, and filopodia formation. Sci Rep, 2017. 7(1).

14. Horsthemke, M., et al., Multiple roles of filopodial dynamics in particle capture and phagocytosis and phenotypes of Cdc42 and Myo10 deletion. J Biol Chem, 2017. 292(17): p. 7258–7273.

15. Plantard, L., et al., PtdIns(3,4,5)P<sub>3</sub> is a regulator of myosin-X localization and filopodia formation. J Cell Sci, 2010. 123(Pt 20): p. 3525–3534.

16. Raines, A.N., et al., Headless Myo10 is a negative regulator of full-length Myo10 and inhibits axon outgrowth in cortical neurons. J Biol Chem, 2012. 287(30): p. 24873–24883.

17. Singh, S.K., et al., Melanin transfer in human skin cells is mediated by filopodia--a model for homotypic and heterotypic lysosome-related organelle transfer. FASEB J, 2010. 24(10): p. 3756–3769.

18. Tokuo, H., K. Mabuchi, and M. Ikebe, The motor activity of myosin-X promotes actin fiber convergence at the cell periphery to initiate filopodia formation. J Cell Biol, 2007. 179(2): p. 229–238.

19. Nagy, S., et al., A myosin motor that selects bundled actin for motility. Proc Natl Acad Sci U S A, 2008. 105(28): p. 9616–9620.

20. Ricca, B.L. and R.S. Rock, The stepping pattern of myosin X is adapted for processive motility on bundled actin. Biophys J, 2010. 99(6): p. 1818–1826.

21. Ropars, V., et al., The myosin X motor is optimized for movement on actin bundles. Nat Commun, 2016. 7.

22. Kerber, M.L., et al., A novel form of motility in filopodia revealed by imaging myosin-X at the single-molecule level. Curr Biol, 2009. 19(11): p. 967–973.

23. Hirano, Y., et al., Structural basis of cargo recognition by the myosin-X MyTH4-FERM domain. EMBO J, 2011. 30(13): p. 2734–2747.

24. Miihkinen, M., et al., Myosin-X and talin modulate integrin activity at filopodia tips. Cell Rep, 2021. 36(11).

25. Zhang, H., et al., Myosin-X provides a motor-based link between integrins and the cytoskeleton. Nat Cell Biol, 2004. 6(6): p. 523–531.

26. Albuschies, J. and V. Vogel, The role of filopodia in the recognition of nanotopographies. Sci Rep, 2013. 3.

27. Johnson, H.E., et al., F-actin bundles direct the initiation and orientation of lamellipodia through adhesion-based signaling. J Cell Biol, 2015. 208(4): p. 443–455.

28. Morgan, M.R., et al., Giving off mixed signals--distinct functions of alpha5beta1 and alphavbeta3 integrins in regulating cell behaviour. IUBMB life, 2009. 61(7): p. 731–738.

29. Wong, S., W.H. Guo, and Y.L. Wang, Fibroblasts probe substrate rigidity with filopodia extensions before occupying an area. Proc Natl Acad Sci U S A, 2014. 111(48): p. 17176–17181.

30. Bornschlogl, T., et al., Filopodial retraction force is generated by cortical actin dynamics and controlled by reversible tethering at the tip. Proc Natl Acad Sci U S A, 2013. 110(47): p. 18928–18933.

31. Lagarrigue, F., et al., A RIAM/lamellipodin-talin-integrin complex forms the tip of sticky fingers that guide cell migration. Nat Commun, 2015. 6.

32. Leijnse, N., L.B. Oddershede, and P.M. Bendix, Helical buckling of actin inside filopodia generates traction. Proc Natl Acad Sci U S A, 2015. 112(1): p. 136–141.

33. Romero, S., et al., Filopodium retraction is controlled by adhesion to its tip. J Cell Sci, 2012. 125(Pt 21): p. 4999–5004.

34. Fischer, R.S., et al., Filopodia and focal adhesions: An integrated system driving branching morphogenesis in neuronal pathfinding and angiogenesis. Dev Biol, 2019. 451(1): p. 86–95.

35. Hu, W., B. Wehrle-Haller, and V. Vogel, Maturation of filopodia shaft adhesions is upregulated by local cycles of lamellipodia advancements and retractions. PLoS One, 2014. 9(9).

36. Jacquemet, G., et al., L-type calcium channels regulate filopodia stability and cancer cell invasion downstream of integrin signalling. Nat Commun, 2016. 7.

37. Jacquemet, G., et al., Filopodome Mapping Identifies p130Cas as a Mechanosensitive Regulator of Filopodia Stability. Curr Biol, 2019. 29(2): p. 202–216.e7.

38. Schäfer, C., et al., One step ahead: role of filopodia in adhesion formation during cell migration of keratinocytes. Exp Cell Res, 2009. 315(7): p. 1212–1224.

39. Gallop, J.L., Filopodia and their links with membrane traffic and cell adhesion. Semin Cell Dev Biol, 2020. 102: p. 81–89.

40. Jacquemet, G., H. Hamidi, and J. Ivaska, Filopodia in cell adhesion, 3D migration and cancer cell invasion. Curr Opin Cell Biol, 2015. 36: p. 23–31.

41. Weber, K.L., et al., A microtubule-binding myosin required for nuclear anchoring and spindle assembly. Nature, 2004. 431(7006): p. 325–329.

42. Woolner, S., et al., Myosin-10 and actin filaments are essential for mitotic spindle function. J Cell Biol, 2008. 182(1): p. 77–88.

43. Kwon, M., et al., Direct Microtubule-Binding by Myosin-10 Orients Centrosomes toward Retraction Fibers and Subcortical Actin Clouds. Dev Cell, 2015. 34(3): p. 323–337.

44. Kwon, M., et al., Mechanisms to suppress multipolar divisions in cancer cells with extra centrosomes. Genes Dev, 2008. 22(16): p. 2189–2203.

45. Varadarajan, R. and N.M. Rusan, Bridging centrioles and PCM in proper space and time. Essays Biochem, 2018. 62(6): p. 793–801.

46. Vasquez-Limeta, A. and J. Loncarek, Human centrosome organization and function in interphase and mitosis. Semin Cell Dev Biol, 2021. 117: p. 30–41.

47. Maiato, H. and E. Logarinho, Mitotic spindle multipolarity without centrosome amplification. Nat Cell Biol, 2014. 16(5): p. 386–394.

48. Alfaro-Aco, R. and S. Petry, How TPX2 helps microtubules branch out. Cell Cycle 2017. 16(17): p. 1560–1561.

49. Petry, S., et al., Branching microtubule nucleation in Xenopus egg extracts mediated by augmin and TPX2. Cell, 2013. 152(4): p. 768–777.

50. Wadsworth, P., TPX2. Curr Biol, 2015. 25(24): p. R1156–R1158.

51. Toyoshima, F. and E. Nishida, Integrin-mediated adhesion orients the spindle parallel to the substratum in an EB1-and myosin X-dependent manner. EMBO J, 2007. 26(6): p. 1487–1498.

52. Beskow, L.M., Lessons from HeLa Cells: The Ethics and Policy of Biospecimens. Annu Rev Genomics Hum Genet, 2016. 17: p. 395–417.

53. Dickinson, M.E., et al., High-throughput discovery of novel developmental phenotypes. Nature, 2016. 537(7621): p. 508–514.

54. Cramer, L. and T.J. Mitchison, Moving and stationary actin filaments are involved in spreading of postmitotic PtK2 cells. J Cell Biol, 1993. 122(4): p. 833–843.

55. Cramer, L.P. and T.J. Mitchison, Myosin is involved in postmitotic cell spreading. J Cell Biol, 1995. 131(1): p. 179–189.

56. Cramer, L.P. and T.J. Mitchison, Investigation of the mechanism of retraction of the cell margin and rearward flow of nodules during mitotic cell rounding. Mol. Cell Biol., 1997. 8(1): p. 109–119.

57. Mitchison, T.J., Actin based motility on retraction fibers in mitotic PtK2 cells. Cell Motil Cytoskeleton, 1992. 22(2): p. 135–151.

58. Taubenberger, A.V., B. Baum, and H.K. Matthews, The Mechanics of Mitotic Cell Rounding. Front Cell Dev Biol, 2020. 8.

59. Sandquist, J.C., M.E. Larson, and K.J. Hine, Myosin-10 independently influences mitotic spindle structure and mitotic progression. Cytoskeleton 2016. 73(7): p. 351–364.

60. Sandquist, J.C., et al., An interaction between myosin-10 and the cell cycle regulator Wee1 links spindle dynamics to mitotic progression in epithelia. J Cell Biol, 2018. 217(3): p. 849–859.

61. Cosenza, M.R. and A. Krämer, Centrosome amplification, chromosomal instability and cancer: mechanistic, clinical and therapeutic issues. Chromosome Res, 2016. 24(1): p. 105–126.

62. Godinho, S.A., M. Kwon, and D. Pellman, Centrosomes and cancer: how cancer cells divide with too many centrosomes. Cancer Metastasis Rev, 2009. 28(1-2): p. 85–98.

63. Rhys, A.D. and S.A. Godinho, Dividing with Extra Centrosomes: A Double Edged Sword for Cancer Cells. Adv Exp Med Biol, 2017. 1002: p. 47–67.

64. Mittasch, M., et al., Regulated changes in material properties underlie centrosome disassembly during mitotic exit. J Cell Biol, 2020. 219(4).

65. Woodruff, J.B., The material state of centrosomes: lattice, liquid, or gel? Curr Opin Struct Biol, 2021. 66: p. 139–147.

66. Kotak, S. and P. Gönczy, NuMA phosphorylation dictates dynein-dependent spindle positioning. Cell Cycle, 2014. 13(2): p. 177–178.

67. Kiyomitsu, T. and S. Boerner, The Nuclear Mitotic Apparatus (NuMA) Protein: A Key Player for Nuclear Formation, Spindle Assembly, and Spindle Positioning. Front Cell Dev Biol, 2021. 9.

68. Kotak, S., Mechanisms of Spindle Positioning: Lessons from Worms and Mammalian Cells. Biomolecules, 2019. 9(2).

69. Seldin, L., et al., NuMA localization, stability, and function in spindle orientation involve 4.1 and Cdk1 interactions. Mol Biol Cell, 2013. 24(23): p. 3651–3662.

70. Long, A.F., P. Suresh, and S. Dumont, Individual kinetochore-fibers locally dissipate force to maintain robust mammalian spindle structure. J Cell Biol, 2020. 219(8).

71. Dix, C.L., et al., The Role of Mitotic Cell-Substrate Adhesion Re-modeling in Animal Cell Division. Dev Cell, 2018. 45(1): p. 132–145.e3.

72. Fink, J., et al., External forces control mitotic spindle positioning. Nat Cell Biol, 2011. 13(7): p. 771–778.

73. Théry, M. and M. Bornens, Cell shape and cell division. Curr Opin Cell Biol, 2006. 18(6): p. 648–657.

74. Théry, M., et al., Experimental and theoretical study of mitotic spindle orientation. Nature, 2007. 447(7143): p. 493–496.

75. Watts, C.A., et al., Design, synthesis, and biological evaluation of an allosteric inhibitor of HSET that targets cancer cells with supernumerary centrosomes. Chem Biol, 2013. 20(11): p. 1399–1410.

76. Tokuo, H., J. Bhawan, and L.M. Coluccio, Myosin X is required for efficient melanoblast migration and melanoma initiation and metastasis. Sci Rep, 2018. 8(1).

77. Kenchappa, R.S., et al., Myosin 10 Regulates Invasion, Mitosis, and Metabolic Signaling in Glioblastoma. iScience, 2020. 23(12).

78. Pozo, F.M., et al., MYO10 drives genomic instability and inflammation in cancer. Sci Adv, 2021. s7(38).

79. Courson, D.S. and R.E. Cheney, Myosin-X and disease. Exp Cell Res, 2015. 334(1): p. 10–15.

80. Hamidi, H. and J. Ivaska, Every step of the way: integrins in cancer progression and metastasis. Nat Rev Cancer, 2018. 18(9): p. 533–548.

81. Bergstralh, D.T., N.S. Dawney, and D. St Johnston, Spindle orientation: a question of complex positioning. Development, 2017. 144(7): p. 1137–1145.

82. Bergstralh, D.T., T. Haack, and D. St Johnston, Epithelial polarity and spindle orientation: intersecting pathways. Philos Trans R Soc Lond B Biol Sci, 2013. 368(1629).

83. Lechler, T. and M. Mapelli, Spindle positioning and its impact on vertebrate tissue architecture and cell fate. Nat Rev Mol Cell Biol, 2021. 22(10): p. 691–708.

84. Pietro, F., A. Echard, and X. Morin, Regulation of mitotic spindle orientation: an integrated view. EMBO Rep, 2016. 17(8): p. 1106–1130.

85. Seldin, L. and I. Macara, Epithelial spindle orientation diversities and uncertainties: recent developments and lingering questions. F1000 Res, 2017. 6: p. 984.

86. Liu, K.C., et al., Myosin-X functions in polarized epithelial cells. Mol Biol Cell, 2012. 23(9): p. 1675–1687.

87. Bockholt, S.M. and K. Burridge, An examination of focal adhesion formation and tyrosine phosphorylation in fibroblasts isolated from src-, fyn-, and yes-mice. Cell Adhes Commun, 1995. 3(2): p. 91–100.

88. Ran, F.A., et al., Genome engineering using the CRISPR-Cas9 system. Nat Protoc, 2013. 8(11): p. 2281–2308.

89. Murugesan, S., et al., Formin-generated actomyosin arcs propel T cell receptor microcluster movement at the immune synapse. J Cell Biol, 2016. 215(3): p. 383–399.

