## Supplementary figures and images for "Myosin 10 supports mitotic spindle bipolarity by promoting PCM integrity and supernumerary centrosome clustering"

### FigS1

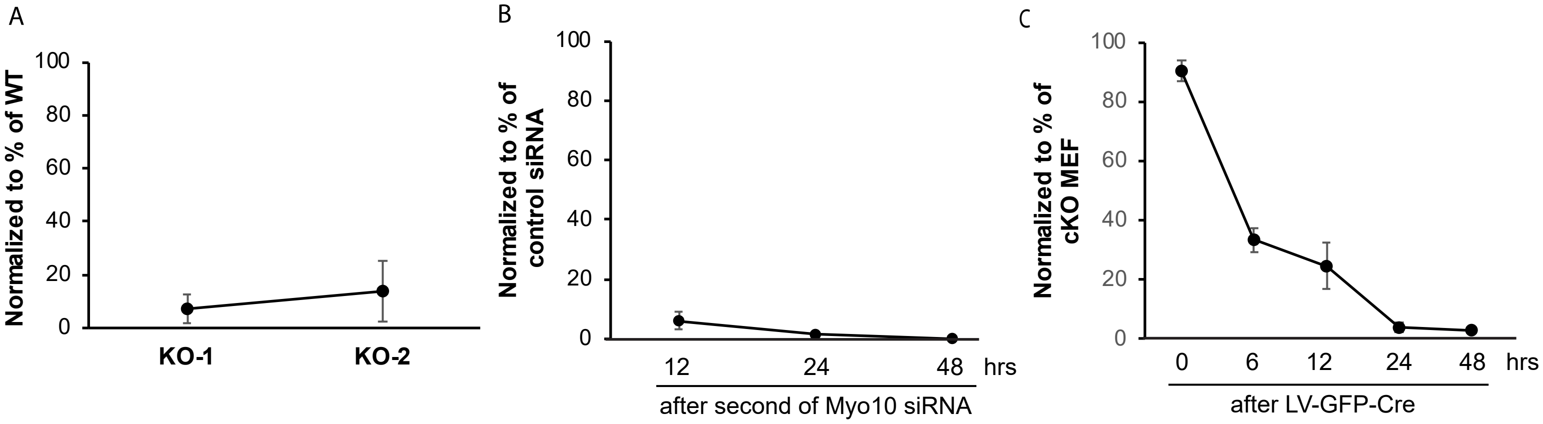

### FigS2

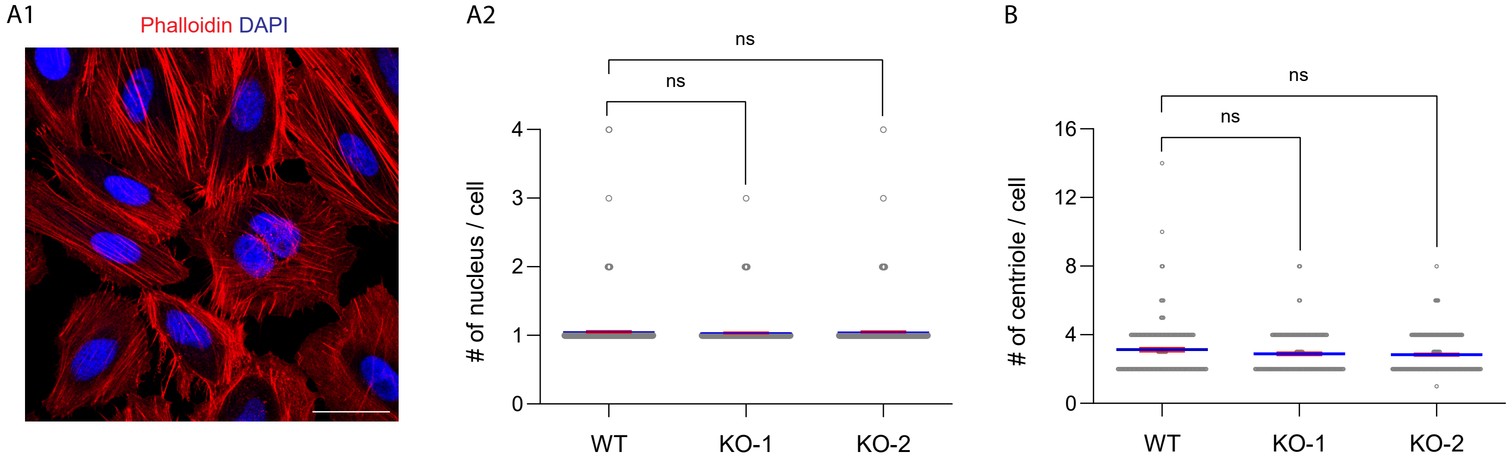

### FigS3

FigS3

A

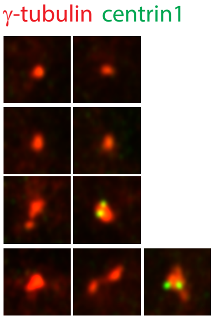


B

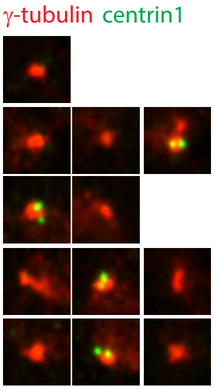
